# Decoding the molecular cues regulating the transition of influenza virus polymerase from transcriptase to replicase

**DOI:** 10.64898/2026.11.05.764532

**Authors:** Aratrika De, Arunava Roy, Arindam Mondal

**Affiliations:** Department of Bioscience and Biotechnology, Indian Institute of Technology (IIT) Kharagpur, Kharagpur, West Bengal, India; Department of Molecular Medicine, University of South Florida, Tampa, Florida, United States of America

**Keywords:** Influenza virus RdRp, polymerase conformation, ANP32, host adaptation, transcription-replication balance

## Abstract

Influenza virus polymerase (FluPol) dynamically transitions between distinct conformational states to coordinate viral gene transcription and genome replication. While vRNP-associated monomeric FluPol supports transcription, host ANP32-dependent asymmetric and ANP32-independent symmetric polymerase dimers mediate genome and antigenome (vRNA/cRNA) replication. How FluPol coordinates between these distinct functional states to maintain the transcription–replication balance remains elusive. Here, we identify novel interdomain interactions within PB2 that govern the stability of the replication-competent conformation of FluPol. A highly conserved salt-bridge between PB2 Mid-E282 and NLS-K752 stabilizes the replicase conformation, while the residue 627 (E or K) dictates ANP32 engagement, together coordinating polymerase dimerization and replication. Notably, bat influenza A viruses H17N10 and H18N11 harbor unconventional E282S and E/K627S substitutions that impair polymerase dimerization, ANP32 interaction, and vRNA/cRNA replication. Introducing these bat-signature substitutions (E282S/K627S) into human H1N1 polymerase results in phosphorylation at both positions in human cells. These findings suggest that phosphorylation may partially compensate for the loss of negative charge at these residues, revealing a potential phosphomimetic adaptation mechanism orchestrating polymerase conformational dynamics. Collectively, our study establishes a conserved PB2 interdomain interaction as a key regulator of FluPol conformational transitions and transcription–replication balance, highlighting a potential role of phosphomimetic adaptations in influenza virus evolution.

## Introduction

Avian influenza A viruses (IAV) are highly adaptable RNA viruses that continually evolve to expand their host repertoire, causing seasonal epidemics and occasional pandemics^1^. Along with IAV, influenza B viruses (IBV) contributes to the seasonal influenza outbreaks while influenza C viruses (ICV) causes milder infection to human^2^. Recent spillover of highly pathogenic H5N1 viruses into cattle and other mammals has heightened concerns over their potential to acquire sustained human-to-human transmission^3^. Hence, understanding the virus evolution trajectory and elucidating the fundamental concepts of virus replication is of utmost importance for developing novel broad-spectrum anti-influenza drugs^4^.

Influenza virus RNA polymerase (FluPol) is the key determinant of influenza virus replication fitness, host tropism and pathogenicity^5–8^. This ∼260-kDa heterotrimeric FluPol comprises the PB1, PB2, and PA subunits and associates with each of the eight negative-sense genomic RNA segments (vRNAs), together with oligomeric nucleoprotein (NP), to form viral ribonucleoprotein complexes (vRNPs)^9–11^. Following entry and uncoating, incoming vRNPs are transported to the nucleus, where FluPol directs both transcription and genome replication^12^. During transcription, the resident FluPol initiates RNA synthesis from the 3′ end of vRNA using a capped primer snatched from the host pre-mRNA^13,14^. The PB2 cap binding domain (CBD) binds the cap and the PA endonuclease domain (PA-endo) cleaves the mRNA to generate a short capped RNA primer, which is subsequently extended by PB1^15,16,14^. Genome replication proceeds through two sequential steps: synthesis of a full-length complementary RNA (cRNA), followed by its use as a template for progeny vRNA synthesis^17,18^. Unlike transcription, replication is initiated by *de novo* synthesis pppApG dinucleotide at the 3’ termini of the vRNA template. For the cRNA template, initiation occurs internally at 4^th^ and 5^th^ nucleotide from the 3’end followed by realignment of the pppApG to the terminal nucleotide prior to elongation^19,20^. The nascent uncapped vRNA/ cRNA strand is encapsidated by a second copy of the FluPol and a pool of monomeric NP, thus producing vRNP/ cRNPs^21–23,18^.

FluPol shows distinct structural organizations in transcription and replication competent states (Supplementary Fig. S1)^24,25^. The central region of the polymerase is composed of the PB1, PB2 N-terminal and PA C-terminal domains which forms a compact catalytic core^25,26^. In addition, the auxiliary domains consist of PA N-terminal endonuclease domain (PA-Endo) and PB2 C-terminal domain (PB2-CTD) that includes cap binding domain (CBD), Mid-linker domain (Mid-link) and the domain harboring the 627^th^ residue (627-domain) and nuclear localization signal (NLS)^27,28,15,16,29,24^. In the transcriptase structure (FluPol_T_), the PB2 CBD is exposed outwards to bind the host mRNA cap while the PA-Endo is also solvent exposed and faces the PB2 CBD at distance ∼50Å (Supplementary Fig. 1a). This ensures cleavage of the mRNA 10-14 nt downstream of the cap^30,14,31^. The PB2 627-domain is packed against the catalytic core, yet solvent exposed, while the NLS domain mostly remains flexible with distal disordered regions^27,32^.

Functional transition of FluPol from transcriptase to replicase mode is governed by (i) extensive structural reorganization^24,33^ (ii) dimerization^34^ and (iii) interaction with host acidic nuclear proteins 32 (ANP32)^35–37^. The ANP32 bound asymmetric dimer has been presumed to be the replication platform (Supplementary Fig. 1b) where the vRNA bound resident FluPol executes genome replication (Replicase; FluPol_R_) and the second FluPol performs NP mediated encapsidation of the nascent vRNA/ cRNA strands (Encapsidase; FluPol_E_) as soon as it emerges from the catalytic core of FluPol_R_, thus nucleating vRNP/ cRNP assembly^35,38,36,37^. In FluPol_R_, the auxiliary domains undergo radical structural reorganization compared with FluPol_T_; the PB2 627-NLS and CBD flip around the Mid-link, which acts as a dynamic hinge, resulting in close juxtaposition of the PB2 627-NLS with PA-endo (Supplementary Fig. 1c)^24,33^. This makes the FluPol_R_ transcription inactive but competent for *de novo* initiation of replication. Notably, the structure of FluPol_E_ is distinct from those of FluPol_R_ and FluPol_T_, which makes it competent only for encapsidation^35,36^. In contrast to the ANP32-bridged asymmetric dimer, apo-FluPol spontaneously oligomerize to form symmetric dimers (Supplementary Fig. 1d), which has been implicated in template realignment during cRNA to vRNA production^34,37^. In this FluPolA symmetric dimer, both polymerases adopt a replicase-like conformation (FluPol_R_*) with the PB2-CTD domain organization resembling that of FluPol_R_^34^ (Supplementary Fig. 1e). Summarily, influenza virus polymerase adopts multiple conformations in its different oligomeric states, each with a distinct functional role in viral transcription and replication. Yet, the exact chronological order and the specific molecular cues that dynamically regulate the structural transition of FluPol_T_ to FluPol_R_ and vice versa, thereby maintaining the transcription-replication balance, remains unclear.

Here we delineate specific molecular interactions that cumulatively govern the stability of the FluPol_R_ over FluPol_T_, hence determining transcription-replication balance and the overall fitness of FluPol. We previously reported distinct restriction signatures, including PB2 K/E627S and E282S, in the Bat influenza A virus polymerases (IAVs; H17N10 and H18N11)^39^, which constitute a deeply divergent lineage from classical IAVs (H1–16 and N1–9)^40–42^. Here, we show that H17N10 polymerase is intrinsically attenuated relative to H1N1 polymerase owing to the atypical serine residues at the PB2-282^nd^ and 627^th^ positions. Notably, H1N1 viruses harboring PB2-E282S and K627S mutations are specifically phosphorylated at these sites during infection, yet replicated more slowly compared to the WT H1N1 virus, suggesting that phosphomimetic adaptations (phospho-S to E) may have contributed to the replication fitness of classical IAVs. Mechanistically, H17N10 polymerase displays robust transcription but reduced replication efficiency, driven by PB2-S282-mediated destabilization of polymerase dimerization and PB2-S627-mediated impairment of its interaction with ANP32A. We further identify a salt-bridge interaction between PB2 E282 and K752 residues linking the Mid-link and 627-NLS domains that selectively stabilizes the FluPol_R_ over the FluPol_T_ conformation. Loss of this interaction in bat FluPol, due to the PB2-E282S substitution, destabilizes the FluPol_R_ conformation, thereby blocking the transition from transcription to replication. Together, these findings reveal a Mid-link–627-NLS molecular cross-talk that regulates FluPol transition from transcriptase to replicase mode and suggest a phosphomimetic evolutionary mechanism underlying the functional diversification of influenza polymerases.

## Results

### PB2 Phosphomimetic Signatures Regulate Polymerase Fitness and Viral Growth

Replication fitness of bat FluPol in non-bat cells remains poorly characterized. Comparative phylogenetic analysis of PB2, PB1, and PA proteins from classical IAVs isolated from diverse hosts revealed a tightly clustered classical IAV lineage, whereas bat IAV polymerases formed a deeply divergent clade with exceptionally long branches, indicating prolonged independent evolution (Supplementary Figs. 2a-c). To determine whether this divergence affects polymerase function, we reconstituted reporter RNPs of human-adapted A/H1N1/WSN/1933 and bat-derived A/H17N10/Guatemala/164/2009 influenza viruses in HEK293T cells using luciferase-based vRNA or cRNA reporter templates together with PB1, PB2, PA, and NP proteins, as described previously^39^. These reporter RNPs either perform transcription to directly produce reporter mRNA from a vRNA template or perform replication to produce more vRNAs through the cRNA intermediate, thus indirectly contributing to the reporter signal (Fig. 1a). The H17N10 polymerase displayed markedly reduced activity compared with H1N1 on both homologous and heterologous promoters (Fig. 1b). Chimeric RNPs generated by swapping individual polymerase subunits further revealed limited functional compatibility between the two polymerases, as substitution of PB1, PB2, or PA substantially impaired activity (Figs. 1c, d). Notably, replacing H1N1 PB2 with that of H17N10 imposed the least functional penalty, suggesting relatively greater compatibility of PB2 between these two polymerases (Fig. 1d). This also indicated that, despite their divergent evolution, bat IAV PB2 retains closer functional relatedness to classical IAVs.

**Figure 1.**
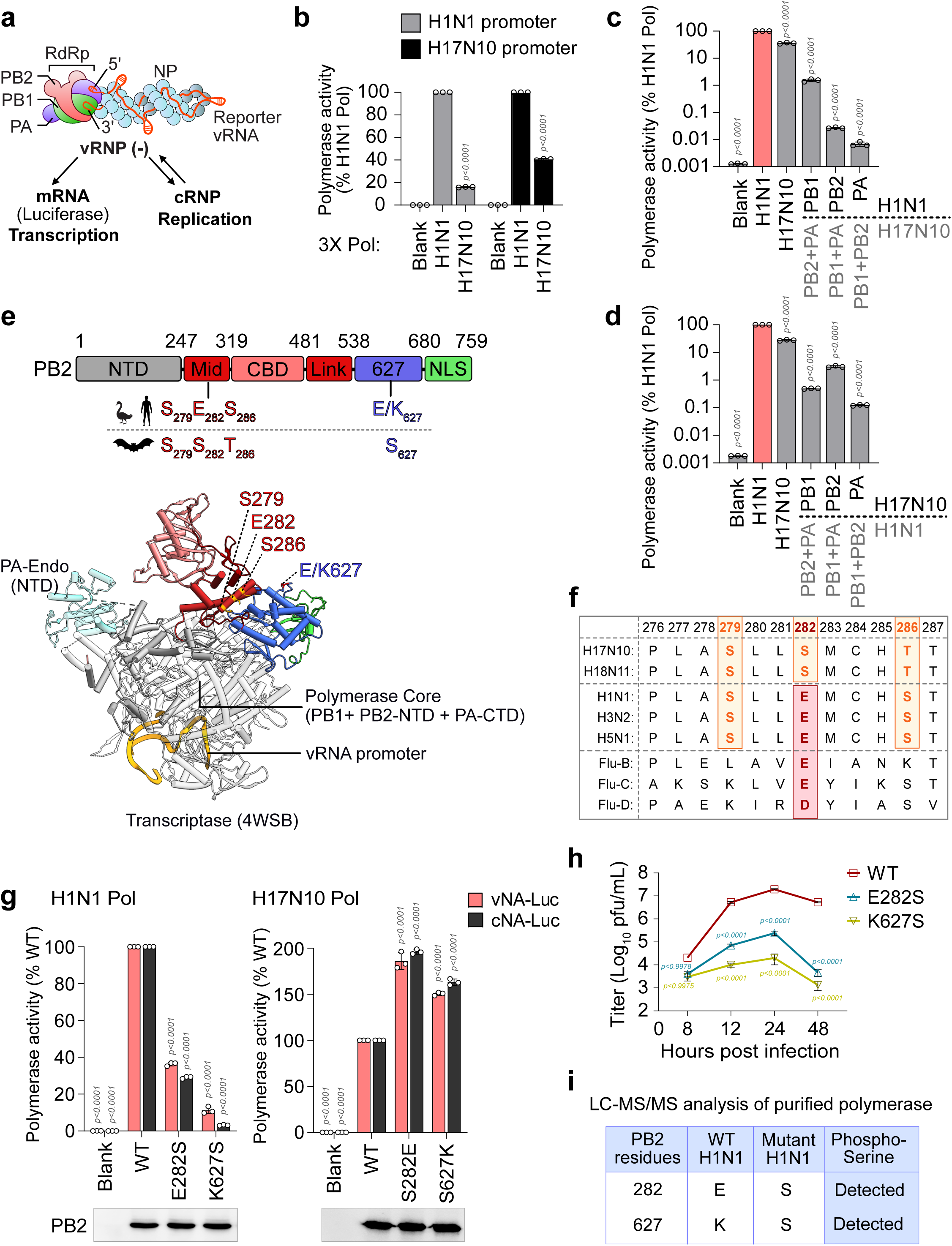
Phosphomimetic signatures in PB2. (a) Schematic of influenza virus RNP and modes of RNA synthesis (adapted from Fodor et al^17^). (b) Comparative polymerase activity of A/H1N1/WSN/1933 and A/H17N10/Guatemala/164/2009 polymerases using homo- or heterotypic promoters. (C–D) Activity of chimeric H17N10 (c) and H1N1 (d) polymerases containing reciprocal subunit swaps, normalized to the respective native polymerases. (e) PB2 domain organization highlighting bat-specific serine residues at positions 282 and 627 and their spatial arrangement in the monomeric transcriptase structure (PDB 4WSB). (f) Conservation analysis of the “S279-E282-S286” motif and E282 residue across influenza A– D viruses. (g) Effect of bat-specific E282S and K627S substitutions on H1N1 polymerase activity and reciprocal substitutions in H17N10 on vRNA and cRNA-sense promoters. (h) Multicycle replication kinetics of H1N1/WSN/1933 viruses carrying WT PB2, PB2-E282S, or PB2-K627S. (i) Summary of PB2-serine phosphorylation in FluPol purified from A549 cells infected with the viruses in (h), as determined by LC-MS/MS. Data are mean ± SD of three biological replicates. P values were determined by one-way ANOVA relative to the wild-type control.

We focused on PB2 to identify molecular signatures underlying the restricted fitness of H17N10 polymerase in human cells. IAV adaptation is strongly influenced by the PB2-627^th^ residue, where avian viruses encode a glutamic acid and human-adapted strains mostly encode a lysine^5,6^. Bat influenza viruses uniquely encode a serine at the 627^th^ position (Fig. 1e). We previously identified another host-adaptive determinant at the 282^nd^ residue within a highly conserved PB2 Mid-domain “S279-E282-S286” motif (henceforth denoted as “S-E-S”) retained across all classical IAVs ^39^. Other influenza virus types also harbor a highly conserved negatively charged glutamate (B and C viruses) or aspartate (D virus) residue at the PB2-282^nd^ position. In contrast, bat influenza viruses (H17N10 and H18N11) encode an atypical “S279-S282-T286” motif (henceforth denoted as “S-S-T”) devoid of a central acidic residue (Figs. 1 e, f). Introduction of the bat-specific E282S or K627S substitutions into H1N1 markedly reduced polymerase activity on both vRNA and cRNA templates. Conversely, restoring the classical IAV signatures (S282E or S627K) substantially enhanced H17N10 polymerase activity in human cells (Fig. 1g). These findings confirm PB2-282S and PB2-627S as bat IAV-specific restriction signatures that constrain H17N10 polymerase fitness in non-bat cells.

Functional consequences of bat-virus specific restriction signatures were further validated using recombinant A/H1N1/WSN/1933 viruses. Mutant viruses carrying either the PB2-E282S or PB2-K627S substitution formed significantly smaller plaques (Supplementary Fig. 2d) and exhibited a 2-4 log reduction in viral titres during multicycle replication in A549 cells, compared to the WT H1N1 (Fig. 1h). Because serine phosphorylation introduces a negative charge that could potentially compensate for the loss of glutamate, we examined if the serine residues introduced at the PB2-282^nd^ or PB2-627^th^ positions get phosphorylated. LC-MS/MS analysis of polymerase purified from cells infected with the respective mutant viruses detected phosphorylation at PB2-282S and PB2-627S (Fig. 1i, Supplementary Figs. 3 a-b). Thus, the conserved glutamate residues at these positions may represent phosphomimetic adaptation signatures that may have evolved in classical IAVs to optimize polymerase activity and replication fitness.

### PB2-E282 creates a conserved acidic surface patch that is critical for influenza polymerase fitness

We next investigated how the PB2-“S279-E282-S286” motif regulates polymerase activity and viral RNA synthesis. Structural analysis of FluPol in the transcriptase conformation revealed a prominent surface-exposed acidic patch contributed by PB2-E282 in IAV/H1N1 (PDB ID: 7NHX) (Fig. 2a) and by the corresponding PB2-E284 residue in influenza B (PDB ID: 4WSA) (Supplementary Fig. 4a). In contrast, the bat IAV/H17N10 polymerase lacks this acidic patch due to the replacement of the canonical “S-E-S” motif with atypical “S-S-T” motif (PDB ID: 4WSB) (Fig. 2b).

**Figure 2.**
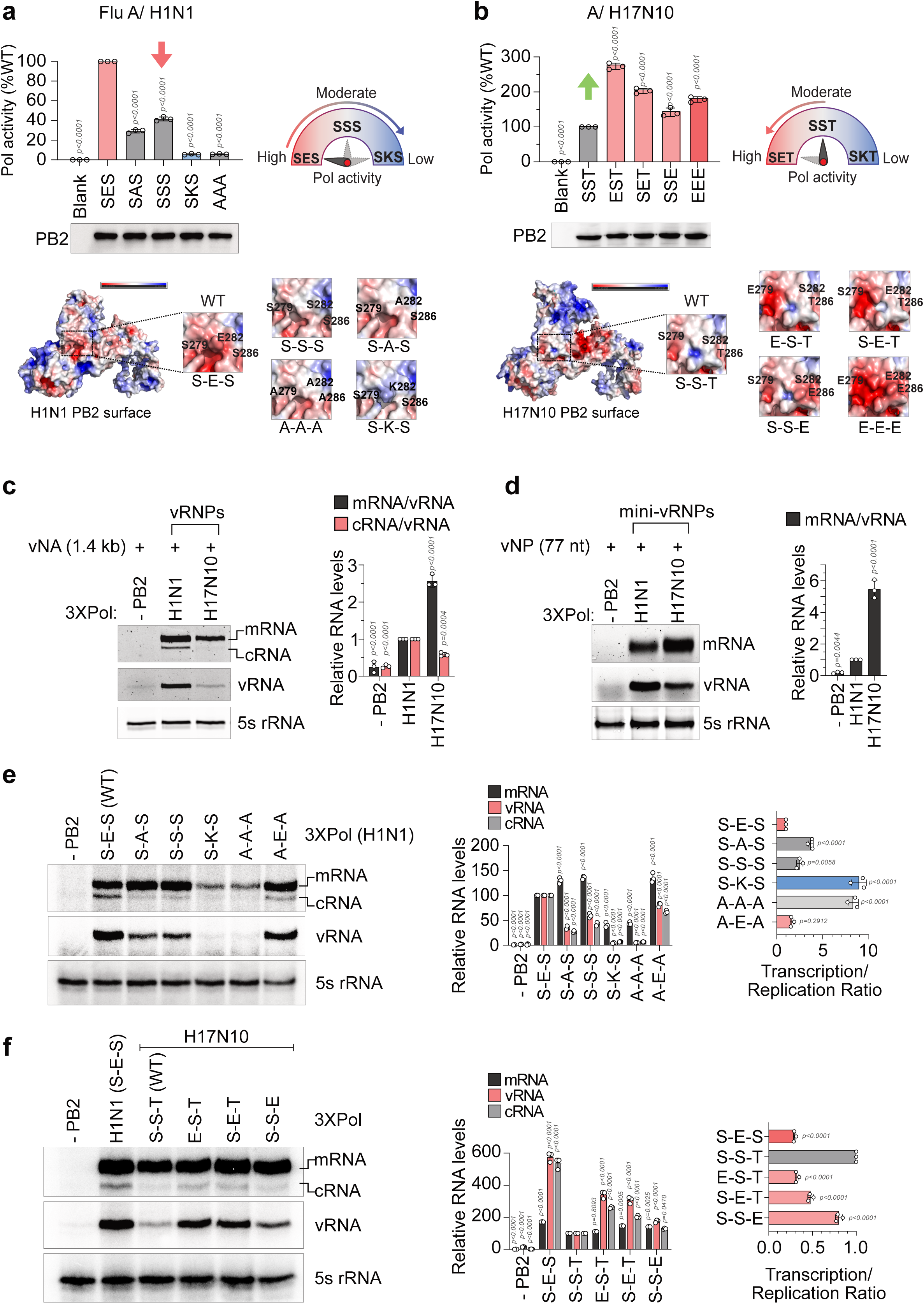
The PB2 E282 residue is critical for maintaining the transcription–replication balance. (a) Polymerase activity of A/H1N1 bearing mutations within the “S279-E282-S286” motif, normalized to WT. (b) Corresponding analysis of A/H17N10 polymerase bearing glutamate substitutions within the native “S279-S282-T286” motif. Schematics indicate the loss or gain of polymerase activity; lower panels show the corresponding PB2 subunits in surface-charge representation (PDB IDs: 7NHX for A/H1N1; 4WSB for A/H17N10); insets highlight mutation-induced changes in surface charge. (c–d) Transcription and replication activities of A/H1N1 and A/H17N10 polymerases either in the context of full-length vRNPs using the vNA template (c) or in the absence of NP using the vNP77 mini-RNA template (d), assessed by primer-extension. Densitometry shows the ratio of positive-sense mRNA/cRNA to vRNA synthesis. (e–f) Transcription and replication activities of WT and mutant A/H1N1 (e) and A/H17N10 (f) polymerases carrying mutations within the respective “S-E-S” or “S-S-T” motifs. Relative abundances of viral transcripts (mRNA) and replicons (c-/vRNAs) are shown, together with the mRNA/vRNA ratio as a measure of the transcription–replication balance.

To determine whether this surface electrostatic feature regulates polymerase activity, we systematically altered the corresponding residues in H1N1, IBV, and H17N10 polymerases. In H1N1, replacing E282 with serine or alanine reduced RNA synthesis by 60% and 70% respectively, whereas introduction of a positively charged lysine or replacement of the entire “S-E-S” motif with triple alanine substitution resulted in ∼95% loss of activity (Fig. 2a). The effect was even more pronounced in IBV, where E284S/A reduced activity by ∼95% and E284K by ∼99% (Supplementary Fig. 4a). Notably, the adjacent IBV PB2-E280 could not compensate for loss of E284, demonstrating that both the presence and precise positioning of the negative charge are critical for optimal polymerase function.

In contrast to H1N1 or IBV, individual alanine substitutions within the H17N10 “S-S-T” motif caused only moderate reductions in activity (30–50%), consistent with minimal alteration of the pre-existing surface electrostatic state (Supplementary Fig. 4b). Likewise, converting the WT “S-S-T” motif to “S-K-T” or “A-A-A” reduced activity by only ∼55–60%, markedly lesser than the corresponding perturbations in H1N1 or IBV (compare Fig. 2a and Supplementary Fig. 4a with Supplementary Fig. 4b). This differential sensitivity towards surface charge alterations is consistent with the already attenuated activity of the H17N10 polymerase, which lacks the acidic surface patch at baseline. We therefore asked whether restoring a negative charge within the “S-S-T” motif could enhance H17N10 polymerase activity. Indeed, introduction of a single glutamate at any position— “E-S-T,” “S-E-T,” or “S-S-E”—increased activity by ∼1.5-to 3-fold relative to wild type (Fig. 2b). However, addition of multiple glutamates (“E-E-T,” “S-E-E,” or “E-E-E”) failed to produce a further combinatorial increase, yielding only a ∼1.5-to 2-fold enhancement (Supplementary Fig. 4c). These findings indicate that FluPol activity depends not simply on the presence of negative charge, but on its precise positioning and optimal density within the PB2 surface patch. Together, these results establish the PB2 acidic surface patch as a conserved electrostatic regulator of influenza polymerase activity.

### The species-specific PB2-282 residue is critical for maintaining the transcription– replication balance of FluPol

Our data suggest that the attenuated activity of the H17N10 polymerase relative to classical IAV polymerases is associated with the bat-specific PB2-E/K627S and E282S substitutions. To identify the affected steps of viral RNA synthesis, we reconstituted H1N1 and H17N10 RNPs as described earlier and performed primer-extension analysis using mRNA-, cRNA- and vRNA-specific probes^43,39^. Compared with H1N1, H17N10 polymerase exhibited markedly reduced cRNA and vRNA synthesis, while mRNA synthesis was largely preserved, indicating a shift in the transcription–replication balance towards transcription (Fig. 2c). This phenotype was recapitulated using NP-free mini-RNPs reconstituted with a 77-nt mini-vRNA template, demonstrating that the altered balance is an intrinsic property of the H17N10 polymerase rather than a consequence of defective RNP assembly (Fig. 2d).

To determine whether the PB2-“S-E-S” motif underlies this phenotype, we introduced targeted substitutions into the H1N1 polymerase. Charge-depleting mutants (“S-S-S” and “S-A-S”) selectively impaired genome replication without substantially affecting transcription, whereas the charge-reversing “S-K-S” mutant and the triple-alanine “A-A-A” mutant caused pronounced vRNA synthesis defects that secondarily reduced transcription. In contrast, the “A-E-A” mutant retained near-WT RNA synthesis, indicating that the central acidic residue at PB2-282 is sufficient to support efficient replication (Fig. 2e). Similarly, substitutions of the corresponding PB2-E284 residue in influenza B virus (FluB) preferentially impaired replication while largely preserving transcription (Supplementary Fig. 5a). Notably, the replication activity of the E282S (for IAV) or E284S (for IBV) mutants remains higher than the corresponding alanine mutants (E282A and E284A), which could be attributed to potential phosphorylation of these serine residues, as already shown for IAV (Fig. 1i).

Introducing glutamate at any position of the H17N10 “S-S-T” motif (“E-S-T”, “S-E-T” or “S-S-E”) largely restored replication, whereas transcription remained unaltered or slightly elevated (Fig. 2f). These findings are consistent with the progressive recovery of RNA synthesis following sequential glutamate substitutions in the H17N10 motif (Fig. 2b), highlighting the importance of the surface-exposed negative charge at PB2-282 in regulating polymerase activity and the transcription–replication balance. Notably, the H1N1 PB2-K627S mutant also exhibited a pronounced replication defect and a comparatively milder transcription defect (Supplementary Fig. 5b), suggesting that this bat-specific substitution similarly shifts polymerase activity towards transcription.

To validate these findings during infection, we quantified viral RNA synthesis in A549 cells infected with WT, PB2-E282S or PB2-K627S viruses. RT-qPCR was performed using by sense-specific primers at different times post-infection as described previously^44^. The E282S virus displayed a moderate transcription defect but a severe replication defect, whereas K627S impaired both processes (Fig. 3a-f). Consequently, the PB2-E282S virus exhibited an increased transcription-to-replication ratio relative to WT, particularly at later time points, while the K627S virus showed transcription-to-replication ratio comparable to that of WT (Fig. 3g). Moreover, the cRNA to vRNA synthesis ratio, reflecting replication efficiency of these viruses, remained attenuated at earlier time-points and only partially recovered later (Fig. 3h). Collectively, these results establish PB2-282 as a key determinant of the transcription– replication balance and indicate that serine substitutions at PB2-282 and PB2-627 contribute to the suboptimal replication activity of the H17N10 polymerase.

**Figure 3.**
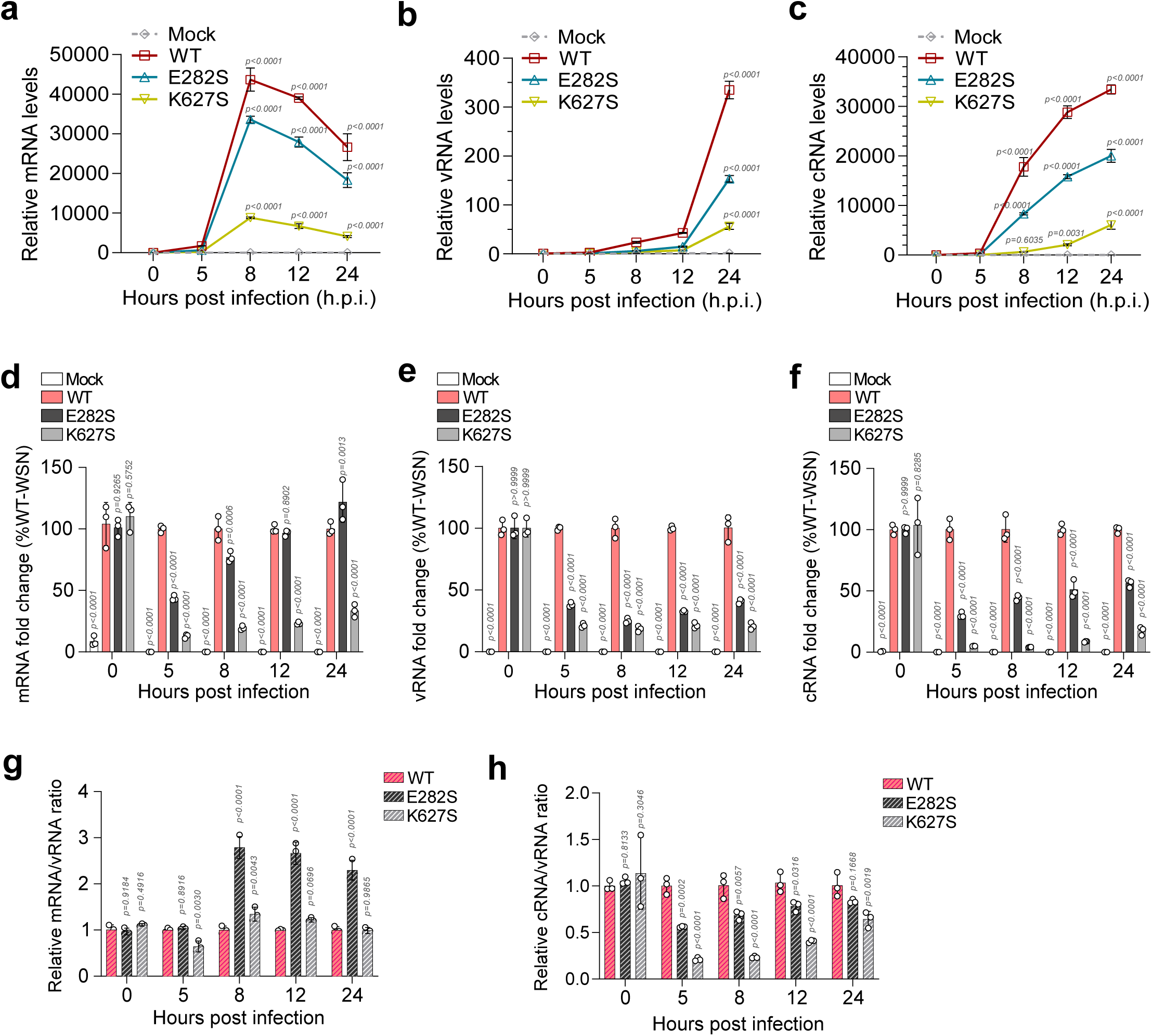
Recombinant A/H1N1 virus harboring H17N10-specific serine at PB2-282 and 627 demonstrates altered transcription–replication balance. Recombinant A/H1N1/WSN viruses harboring the E282S or K627S substitution were used to infect A549 cells, followed by RNA isolation and qRT-PCR to quantify viral RNA levels. Time kinetics of (a) mRNA, (b) vRNA and (c) cRNA synthesis relative to the input levels (0 h.p.i) are represented for WT and mutant viruses. (D-F) Relative abundance of mRNA (d), vRNA (e) and cRNA (f) for mutant viruses at different h.p.i, normalized to the WT virus. mRNA/vRNA (g) and cRNA/vRNA (h) ratios calculated at different h.p.i., representing transcription and replication efficiency, respectively.

### PB2-E282 is critical for efficient initiation of genome replication

Influenza genome replication involves FluPol_R_-mediated RNA synthesis and FluPol_E_-mediated encapsidation of nascent RNA by NP^35,38^. To investigate which of these steps is regulated by PB2-E282, we performed a cRNA stabilization assay using cells pre-expressing NP and a catalytically inactive polymerase (Fig. 4a, left)^21^. Following infection with A/H1N1/WSN and cycloheximide treatment, incoming RNPs supported primary transcription and, in collaboration with the pre-expressed polymerase and NP, supported vRNP-to-cRNP formation. However, newly assembled cRNPs containing catalytically inactive polymerase could not support subsequent cRNA-to-vRNA replication as previously described^45^. Pre-expressed WT and PB2-E282 mutant polymerases supported comparable mRNA and cRNA accumulation under these conditions, indicating that PB2-E282 is dispensable for encapsidation (Fig. 4b). Next, we repeated the same experiment with catalytically active polymerase, which can support consecutive rounds of cRNA and vRNA synthesis through its encapsidase and replicase activities (Fig. 4a, right), as demonstrated earlier^46^. Now, the polymerases harboring PB2-E282 mutations showed markedly reduced cRNA as well as vRNA synthesis compared to the WT (Fig. 4c). Similar results were also observed for the PB2-E284 mutants in FluB polymerase (Supplementary Fig. 6). The defect in genome replication was further exacerbated when incoming viral RNPs were supplied by the mutant PB2-E282S virus, markedly reducing cRNA and vRNA accumulation while increasing mRNA levels (Fig. 4d). These findings implicate defective replication, rather than encapsidation, as the primary consequence of PB2-E282 mutation.

**Figure 4.**
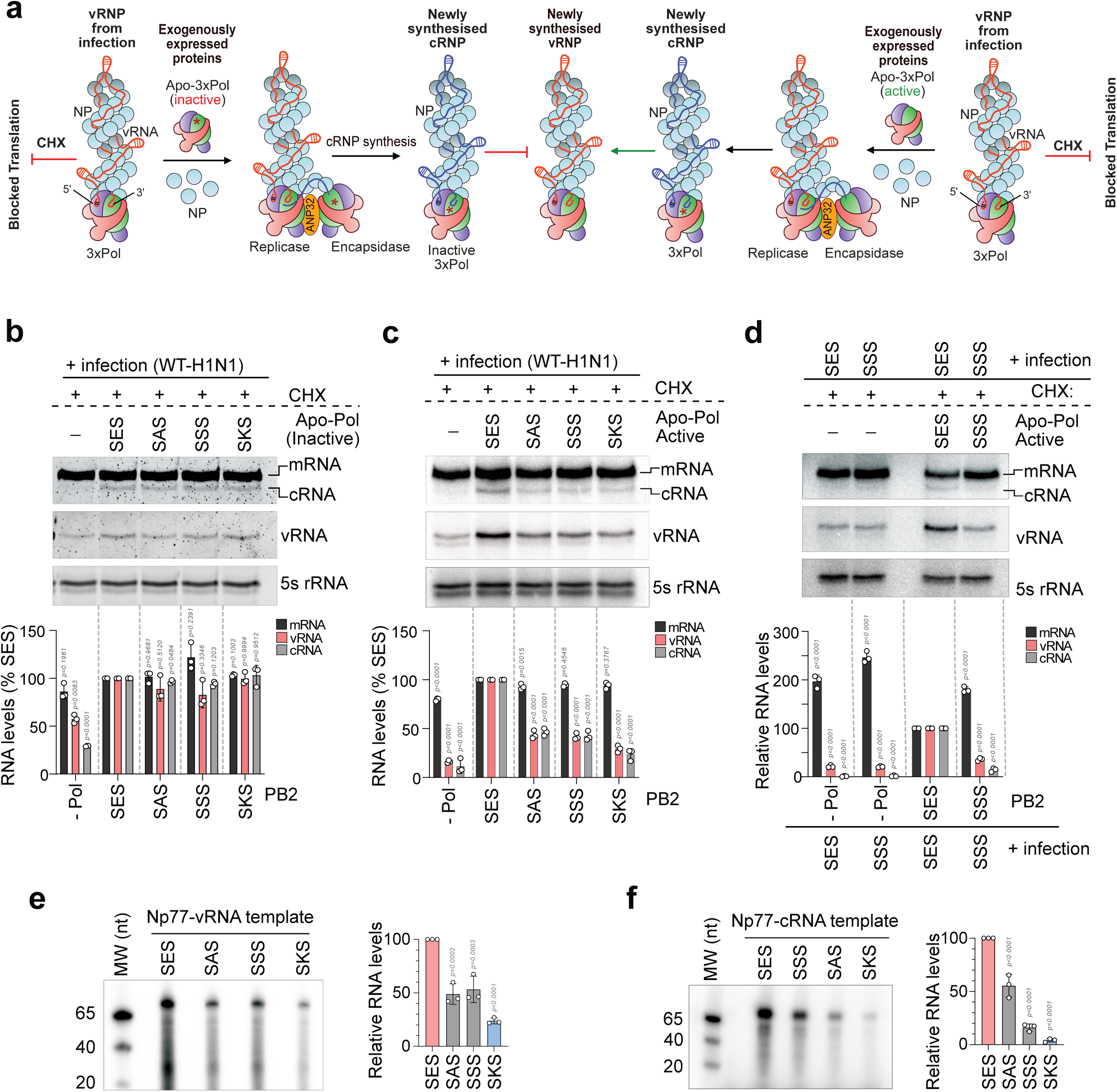
The PB2 E282 residue regulates cRNA/vRNA replication. (a) Schematic of the cRNA/vRNA stabilization and replication assay. Cells pre-expressing NP and either catalytically active or inactive (PB1-A: 445/446A mutant) polymerases containing WT or “S-E-S” motif-mutant PB2 were infected with WT A/H1N1/WSN virus and treated with cycloheximide at 0 h.p.i. (b–c) Viral RNA synthesis measured by primer extension. Catalytically inactive polymerase supported mRNA and cRNA, but not vRNA synthesis (b), whereas catalytically active polymerase supported the synthesis of mRNA, cRNA, and vRNA (c). (d) Cells pre-expressing NP and catalytically active polymerases containing either WT (E282) or E282S mutant PB2 were infected with WT A/H1N1/WSN or PB2-E282S mutant viruses, followed by cycloheximide treatment at 0 h.p.i.; viral RNA synthesis was assessed by primer extension. Relative abundances of viral transcripts (mRNA) and replicons (c-/vRNAs) are shown below the corresponding images. (e–f) *In vitro* replication initiation assays using purified WT or “S-E-S” motif-mutant polymerases with negative-sense vNP77 (e) or positive-sense cNP77 (f) RNA templates in the presence of ApG and rNTPs, analyzed by denaturing PAGE.

To directly assess the role of PB2-E282 in replication, we reconstituted an *in vitro de novo* replication assay using 77-nt vRNA or cRNA templates, an ApG dinucleotide primer, rNTPs, and [α-^32^P]-UTP^19,43,47^. WT polymerase efficiently replicated the mini-v/c-RNA templates in the absence of NP, whereas the PB2-E282 mutants exhibited moderate-to-severe reductions in replication from both templates, correlating with the extent of charge depletion (Figs. 4 e, f). Collectively, these results establish the PB2-E282 residue, located in the Mid-domain, as a critical determinant of efficient FluPol replicase function by specifically regulating the initiation of genome replication.

### The PB2 Mid-domain allosterically regulates FluPol symmetric dimerization

FluPol dimerization is a pre-requisite for its replication activity. While the ANP32-bridged asymmetric dimer is instrumental in vRNA-to-cRNA synthesis and vice versa, ANP32-independent symmetric dimerization is presumed to be critical for cRNA-to-vRNA synthesis^34,35,18^. We therefore investigated whether PB2-E282 regulates polymerase replication activity through regulation of either mode of dimerization. Structural analysis revealed that PB2-E282 lies distal to both dimer interfaces, whereas PB2-K627 is positioned near the asymmetric interface and contributes to ANP32 engagement (Figs. 5a, i).

**Figure 5.**
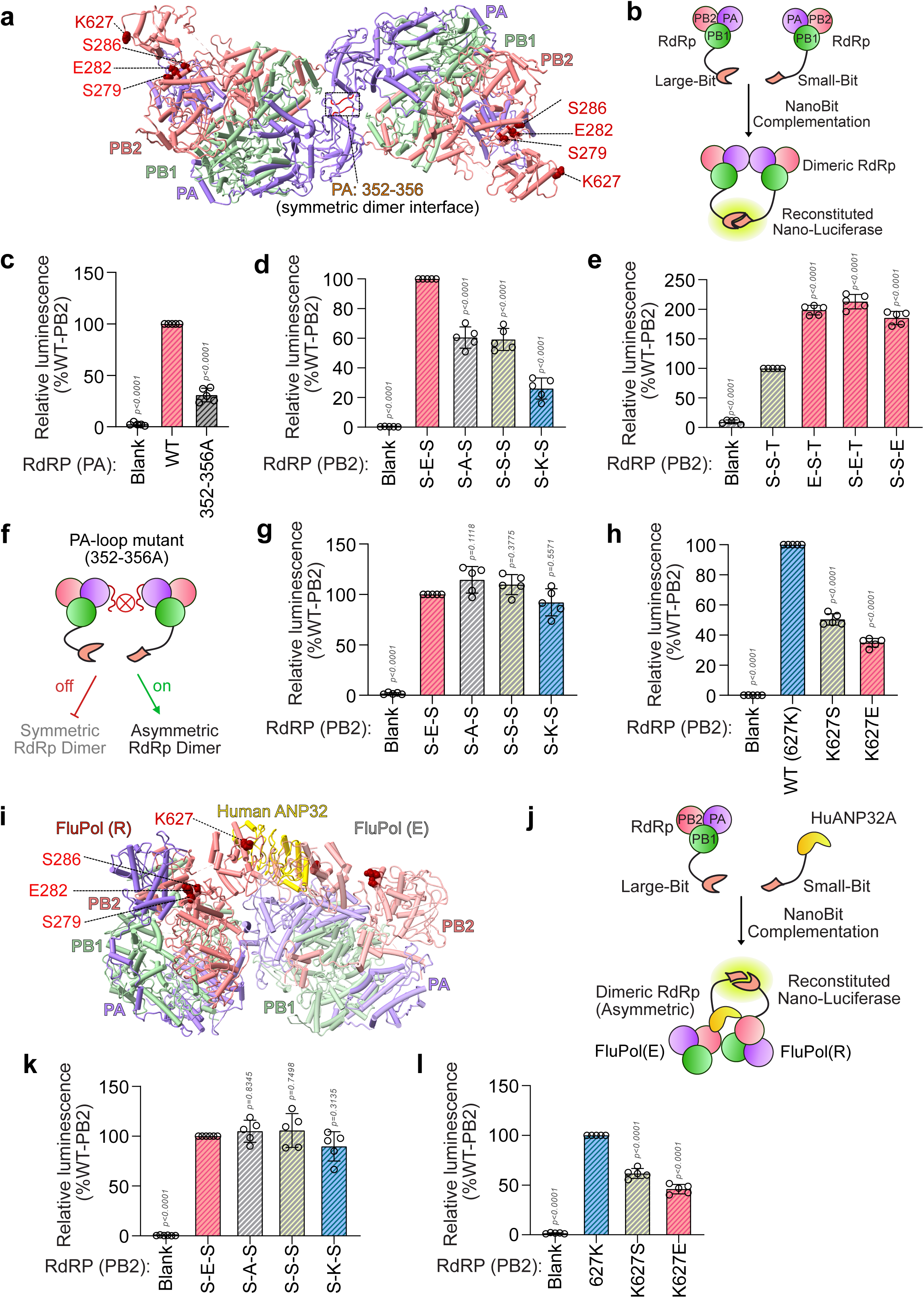
PB2 E282 and K627 regulate polymerase dimerization and ANP32 interaction. (a) Structure of the symmetric FluPol dimer, with PB1, PB2, and PA shown in green, pink, and purple, respectively (PDB ID 6QPF). The PB2-“S279-E282-S286” motif and K627 are highlighted as red spheres, and the PA 352–356 loop forming the dimer interface is shown in red. (b) Schematic of the NanoBiT complementation assay for monitoring FluPol dimerization. (c) Reduced dimerization of the symmetric dimerization-interface mutant PA 352–356A compared with WT. (d–e) Relative dimerization efficiency of A/H1N1 (d) and A/H17N10 (e) polymerases carrying mutations within the respective PB2-“S279-E282-S286” and “S279-S282-T286” motifs, normalized to WT. (f) Schematic of the NanoBiT assay using the PA 352– 356A mutant to assess asymmetric dimer formation. (g–h) Effects of mutations at PB2 E282 (g) and K627 (h) on asymmetric dimer formation in the PA 352–356A background as shown in (f). (i) Structure of the asymmetric dimer showing heterotrimers in the replicase (FluPol_R_) and encapsidase (FluPol_E_) conformations, with the PB2-“S279-E282-S286” motif and K627 highlighted in red and ANP32 shown in yellow (PDB ID 8R1J). (j) Schematic of the NanoBiT assay for monitoring FluPol–ANP32 interaction. (k–l) Effects of mutations at PB2-E282 (k) and K627 (l) on FluPol interaction with human ANP32A.

To monitor FluPol dimerization, we established a NanoBiT complementation assay using SmBiT-and LgBiT-tagged PB1 subunits (Fig. 5b)^48^. WT H1N1 polymerase generated a robust luminescence signal, consistent with efficient dimer formation. Disrupting the previously characterized symmetric dimer interface using PA-352–356A reduced the signal by approximately 70%, indicating that symmetric dimers account for a substantial fraction of the measured complementation, as also reported previously (Fig. 5c)^34,48^. Notably, PB2-E282 mutations markedly impaired H1N1 polymerase dimerization, with E282K producing the strongest defect (Fig. 5d), consistent with their impaired replication activity (Figs. 4 e, f). Co-immunoprecipitation assays independently confirmed the requirement for PB2-E282 in efficient polymerase dimerization (Supplementary Fig. 7a). Conversely, H17N10 polymerase displayed reduced dimerization relative to H1N1, which was enhanced two- to fourfold by introducing glutamate into the species-specific “S-S-T” motif (Fig. 5e; Supplementary Fig 7b).

Strikingly, the effects of PB2-E282 mutations on dimerization were abolished in the symmetric-interface-defective PA-352–356A background (Figs. 5 f, g; Supplementary Fig. 7c), supporting a specific role for PB2-E282 in regulating symmetric dimer formation. In contrast, PB2-K627S and K627E continued to affect dimerization in this background, consistent with their distinct contribution towards ANP32-mediated asymmetric dimer formation (Fig. 5h). To validate this further, we examined FluPol–ANP32A interactions using NanoBiT complementation between PB1-LgBiT and human ANP32A-SmBiT (Fig. 5j)^48,49^. PB2-E282 substitutions had no effect on FluPol-ANP32A association, whereas K627S and K627E significantly reduced the interaction (Figs. 5 k, l), consistent with previous reports^50^. Co-immunoprecipitation also confirmed the results obtained from the NanoBiT complementation assay (Supplementary Fig. 8).

Collectively, these findings reveal two distinct but complementary determinants of FluPol dimerization: PB2-E282 allosterically regulates symmetric dimer formation, whereas PB2-K627 supports ANP32-mediated asymmetric dimerization. The atypical serine substitutions at these positions in H17N10 are therefore associated with defects in both dimerization modes, providing a mechanistic explanation for its impaired genome replication. Additionally, the data presented here suggest that our NanoBiT complementation assay system is efficient in characterising phenotypes involved in symmetric and asymmetric dimerization of the polymerase.

### Interaction between PB2-Mid and NLS domains stabilizes FluPol replicase over transcriptase

To elucidate how PB2-E282 allosterically regulates symmetric dimerization, we compared published structures of monomeric transcriptase and symmetric and asymmetric replicase complexes^14,34,51,35–37^. Notably, both protomers of the symmetric dimer and one protomer of the asymmetric dimer adopt replicase-like conformations (Supplementary Fig. 1e)^18,34^. We therefore sought conserved interactions involving PB2-E282 that selectively stabilize the replicase conformation. We reasoned that such interaction may stabilize the FluPol in its replicase conformation, which could be a pre-requisite for symmetric dimer formation. Structural analysis of the IAV/H5N1 FluPol replicase revealed a close association between the PB2-NLS and PA endonuclease domains, with the C-terminal α-helix of the NLS positioned in close proximity to the PB2-Mid domain helix containing the conserved “S-E-S” motif (Fig. 6a and Supplementary Fig. 9a). Notably, PB2-K752 within the NLS lies approximately 2.7 Å from PB2-E282, suggesting a potential salt-bridge interaction between these two residues (Fig. 6a, inset WT). Substitutions at E282 could disrupt this interaction: E282S or E282A would eliminate the acidic side chain, whereas E282K would introduce electrostatic repulsion and steric clash with K752 (Fig. 6a, insets and table). Interestingly, K752 resides within the bipartite nuclear localization signal of PB2 and is highly conserved across influenza A (including Bat IAVs), B and C viruses, with arginine at the corresponding position in influenza D virus (Fig. 6f and Supplementary Fig. 10a). Thus, the E282-K752 salt-bridge represents a highly conserved intramolecular interaction across all different influenza viruses (Fig. 6a, Supplementary Figs. 9 b, c). Bat IAVs (H17N10 and H18N10) lack this interaction due to the E282S substitution, which might be the root-cause of their replication defect. Notably, glutamic acid substitutions at positions 279 or 282 (“E-S-T” or “S-E-T” motifs, respectively) are predicted to re-establish salt-bridge interactions with K752, whereas glutamic acid substitution at 286 (“S-S-E” motif) has the potential to form a H-bond with N748, thereby stabilizing the replicase conformation (Supplementary Fig. 9d). This explains the position-dependant gain in replication activity observed in Figs. 2b and 2f.

**Figure 6.**
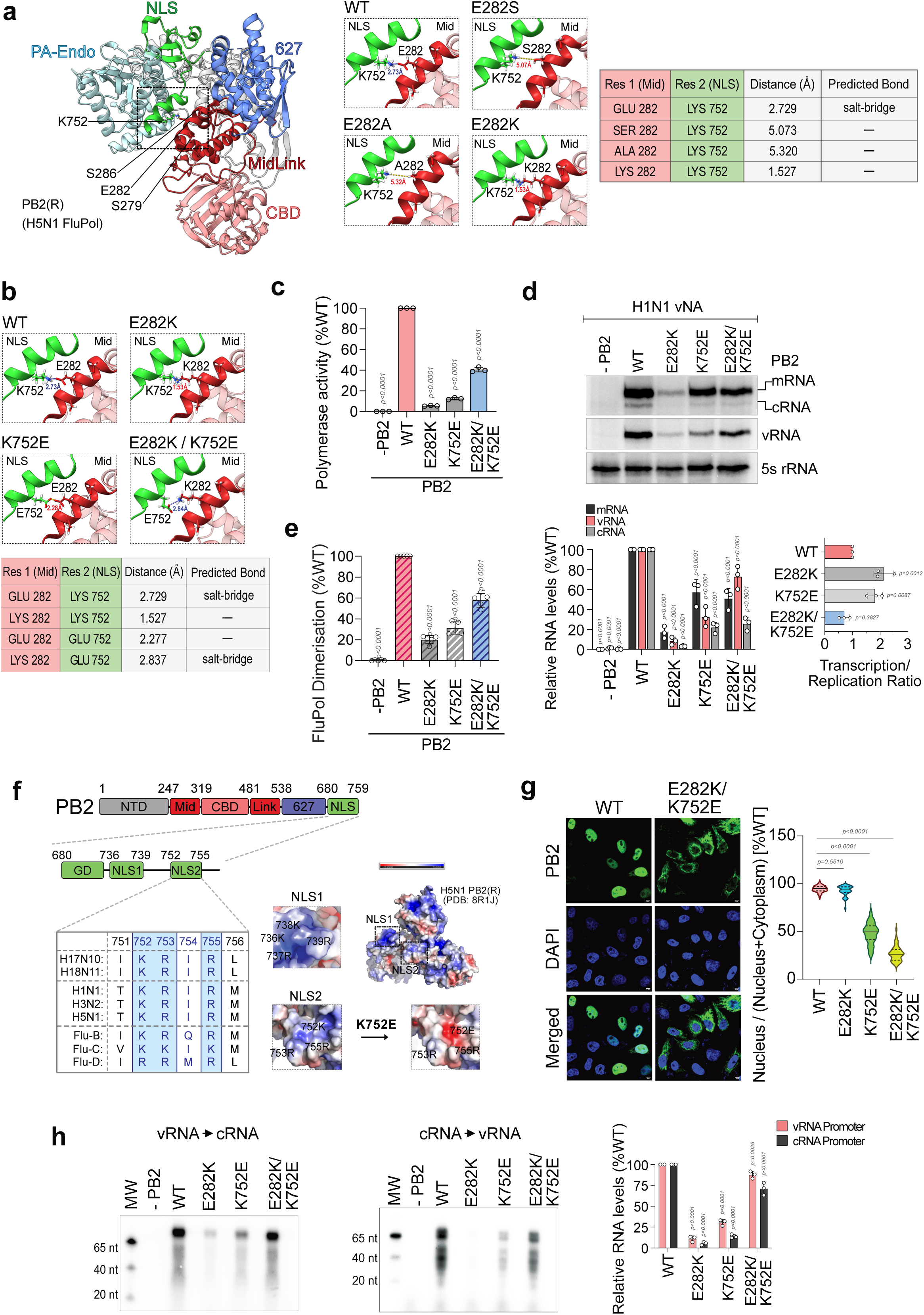
Mid (E282)-NLS (K752) interaction is crucial for driving viral genome replication. (a) Structure highlighting the relative positioning of the PA-endonuclease domain and PB2 subunit in the replicase conformation (FluPol_R_) with different domains represented in different colors; the PB2-“S279-E282-S286” motif in the Mid-domain (red) and K752 in the NLS (green) are shown in stick representation. Insets show the effect of charge-altering mutations at PB2-282; table shows side chain distances and potential interactions between PB2-282 harboring different substitutions and K752 residues, as predicted using UCSF Chimera (b) Structural analysis of charge-swapping mutations at either PB2-282, 752 or both positions, with corresponding side chain distances and potential salt-bridge interactions presented in the table (PDB ID 8R1J). Dotted yellow lines represent interatomic distances, while dotted blue lines indicate potential salt-bridge interactions (c-d) Activity of the polymerase harboring charge-reversal mutations (E to K or vice versa) either at PB2-282, 752 or both positions as analyzed using reporter assay (c) and primer extension analysis; densitometry showing the relative abundance of mRNA, vRNA and cRNAs compared with WT and the ratio between transcription and replication activities are presented (d). (e) The same variants of the polymerase as in (d) are subjected to NanoBiT complementation to assess polymerase dimerization. (f) Schematic of PB2 domain organization, highlighting the NLS-domain with its NLS1 and NLS2 motifs; lower panel shows conservation analysis of 752 residue across influenza A-D viruses; right panel shows surface charge representation of PB2 with positively charged surface patches corresponding to NLS1 and NLS2, and the effect of K752E substitution upon the surface charge of NLS2 (PDB ID 8R1J). (g) Nuclear localization of WT or K752-mutant PB2, as evidenced by immunofluorescence assay and its quantitation. (h) Replication efficiency of the polymerase harboring charge-reversal mutations (E to K or vice versa) either at PB2-282, 752 or both positions on the vRNA or cRNA template, as analyzed by *in vitro* replication assay.

To test this model, we introduced the PB2-K752E mutation into either the WT or E282K polymerase. We speculated that the K752E mutation in WT PB2 would introduce steric clash and electrostatic repulsion with E282, thereby destabilizing the replicase conformation and disrupting polymerase activity similar to the PB2-E282K mutation. In contrast, introducing the K752E mutation in the PB2-E282K polymerase should re-establish the salt-bridge interaction and help the polymerase regain its activity (Fig. 6b). As predicted, the PB2-K752E mutation resulted in drastic 85% reduction in RNA synthesis compared to the WT, whereas introducing K752E into the E282K polymerase exhibited only 50% reduction relative to WT. Thus, the double mutant displayed a 35–45% increase in activity relative to the corresponding single mutants (E282K and K752E), consistent with restoration of the interaction through charge complementation (Fig. 6c). Primer-extension analysis confirmed that these effects primarily reflected altered genome replication, where the single mutants drastically affected c-/v-RNA synthesis, and the double mutants partially restored the same (Fig. 6d). Consistently, NanoBiT complementation revealed impaired dimerization in both PB2 E282K and K752E single mutants and partial restoration of dimerization (∼60%) in the double mutant (Fig. 6e), linking the E282–K752 interaction to efficient dimer formation and genome replication activities of FluPol.

The incomplete rescue of polymerase activity and dimerization by the double mutant suggested that introduction of a charge reversal mutation in the NLS may have other adverse effects that indirectly affect the aforementioned phenotypes. Indeed, K752E disrupts the positively charged surface of the bipartite NLS2 (Fig. 6f), reducing PB2 nuclear localization (Fig. 6g, Supplementary Fig. 10b), and thus should interfere with the assembly of functional RdRp within the nucleus, consistent with previous reports^27,52^. We therefore assessed replication independent of cellular localization using *in vitro de novo* replication assay with mini-vRNA and mini-cRNA templates. Either of the single mutations markedly impaired replication, while the E282K/K752E double mutant restored replication activity to near-WT levels (Fig. 6h). These results support the model in which the PB2-E282–K752 interaction stabilizes the replicase conformation and promotes symmetric dimerization, which is independent of the nuclear localization function of the NLS domain. Disruption of this interaction may therefore contribute to the replication defect of bat influenza polymerases lacking the conserved acidic residue at PB2-282.

### Concentration-dependent symmetric dimerization rescues the replication defect of PB2-E282S polymerase

Finally, we sought to elucidate the potential relationship between the FluPol_T_-to-FluPol_R_ conformational transition and polymerase dimerization, two processes that are regulated by the PB2-E282 residue. We hypothesized that these processes might be coupled and that enhanced symmetric dimer formation could compensate for the impaired ability of the PB2-E282S polymerase to adopt the replicase conformation, thereby rescuing the associated replication defect. To validate this hypothesis, we reconstituted *in vitro* replication using either WT polymerase or increasing concentrations of the defective E282S polymerase. Although the E282S polymerase remained defective relative to WT at equivalent concentrations, increasing its concentration progressively rescued replication, ultimately restoring activity to WT levels (Figs. 7 a, c). In contrast, increasing the concentration of the E282S polymerase carrying the symmetric dimerization-defective PA-352–356A mutation produced only a modest dose-dependent increase in replication and failed to substantially rescue the defect (Figs. 7 b, c). Summarily, the dose dependent increase in the replication activity of the PB2-E282S polymerase becomes much more evident for the dimerization-competent than the dimerization-defective polymerase (Fig. 7 d).

**Figure 7.**
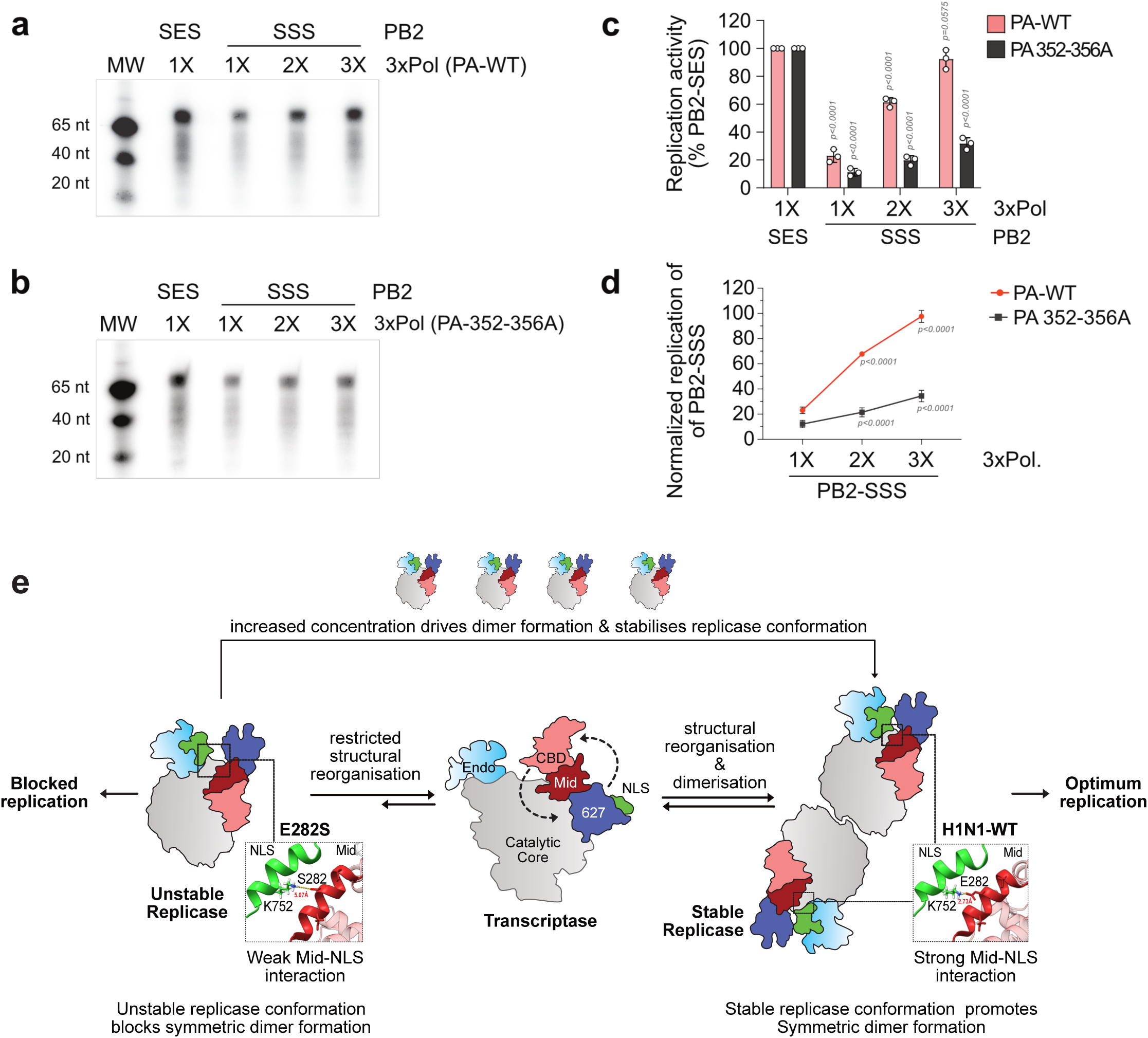
Polymerase symmetric dimerization stabilizes replicase conformation. (a) *In vitro* replication assay reconstituted with either WT polymerase (PB2-“S279-E282-S286”) or increasing concentrations of the mutant polymerase harboring the PB2-“S279-S282-S286” motif. (b) Same experiment as (a) except that the PB2-WT or mutant polymerases were reconstituted with the PA-352-356A mutant. (c-d) Densitometry analyses of (a) and (b) showing either the replication activity of the WT or mutant polymerases (c) or rescue of the replication defect in the context of dimerization-competent (PA-WT) or dimerization-defective (PA-352-356A) PA (d). (e) Proposed model depicting the transition of the transcriptase to a symmetric dimeric replicase stabilized by the PB2-Mid (E282)–NLS (K752) interaction. The E282S mutation disrupts this interaction, thereby destabilizing the replicase conformation, blocking symmetric dimer formation and impairing genome replication. Conversely, increasing concentration of the polymerase can force the conformational switch from transcriptase to replicase through symmetric dimer formation.

Thus, concentration-dependent rescue of the E282S replication defect requires an intact symmetric dimerization interface, linking replicase conformational stability to dimer formation. These findings support the model (Fig. 7e) in which symmetric dimerization promotes the transcriptase-to-replicase transition, while stabilization of the replicase conformation reciprocally facilitates dimer formation. The coupling between these processes is therefore critical for optimal influenza polymerase replication activity.

## Discussion

### Molecular cues defining the transition from FluPol_T_ to FluPolR

Influenza polymerase adopts distinct conformational states to execute transcription and replication, and efficient switching between these states is essential for productive infection^24,33^. Host factors, including RNA Pol II CTD and ANP32A/B, stabilize the transcription and replication-competent states (FluPol_T_ and FluPol_R_), respectively^53,54,37,36^. However, purified FluPol can independently execute both transcription and replication *in vitro*, indicating that the polymerase contains intrinsic molecular determinants that enable its structural recycling between these functional states^55,56^. Here, we identify intra-as well as intermolecular interactions that selectively stabilize the FluPol_R_ conformation, promote FluPol dimerization and thereby play a critical role in optimizing genome replication.

Structurally, FluPol_T_ is characterized by a face-to-face orientation of the PB2-CBD and PA-endo (Supplementary figure 1a), positioned at an optimal distance for cap-snatching^30,14^. In FluPol_R_, the entire PB2-CTD flips, bringing the 627-NLS double domain into close proximity to PA-endo and thereby occluding its catalytic site^24,37,36^. Recent structural studies of H5N1 polymerase suggest that this transition proceeds through an intermediate conformation in which the PB2-CTD undergoes partial rotation before adopting the final replicase conformation^33^. This rearrangement progressively increases the separation between the PB2-CBD and PA-endo domains observed in FluPol_T_. A subsequent large-scale rotation of the PB2-CTD is accompanied by repositioning of the PB2-627 domain and exposure of the ANP32-binding surface. This movement also reorganizes the Mid-domain and brings it into proximity with PA-endo. Concurrently, the previously disordered PB2-NLS becomes structured and packs against the PB2-Mid and PA-endo domains^36,37^.

It is within this replication-competent configuration that K752 in the PB2-NLS approaches the conserved PB2-“S279-E282-S286” motif in the Mid-domain, enabling a potential E282–K752 salt bridge interaction (Fig. 6 a). This interaction is further reinforced by contacts between PB2-NLS and PA-endo, thus providing an intrinsic structural constraint that stabilizes the FluPol_R_ state (Supplementary Fig. 9a). Thus, the E282–K752 interaction may represent a previously unrecognized molecular cue that biases the conformational landscape of FluPol towards replication and establishes a structural platform competent for ANP32 engagement and asymmetric dimer formation.

### Mid-NLS interaction as a critical regulator of FluPol dimerization

Following translation and assembly, apo-FluPol undergoes spontaneous oligomerization, generating dimers and higher-order assemblies^57,34,37^. Recent reports suggests that ANP32 proteins function as electrostatic chaperones that regulate this process and stabilize apo-FluPol in a symmetric dimeric configuration^37^. These ANP32-stabilized symmetric dimers have been proposed to represent a major nuclear reservoir of polymerase, which can subsequently engage resident vRNP-bound polymerase to generate the asymmetric dimeric replication complex. Structural studies indicate that the IAV apo-FluPol symmetric dimers harbor 2-fold symmetry for the core interface while the peripheral domains adopt a replicase-like (FluPol_R_*) conformation^34,37^ (Supplementary Fig. 1e). Our findings place the PB2-Mid–NLS interaction within this regulatory framework. The E282–K752 interaction not only stabilizes the FluPol_R_ conformation but also promotes formation of the symmetric dimer. Disruption of this interaction by E282K or K752E substitutions compromises replication activity and reduces polymerase dimerization (Figs. 6 c-e). Importantly, increasing polymerase concentration, and consequently promoting symmetric dimer formation, substantially rescues the replication defect of the E282S mutant (Figs. 7 a-d). These findings suggest that symmetric dimerization can function as a molecular trigger that reinforces the replication-competent conformation of newly assembled apo-FluPol (Fig. 7e).

This mechanism provides an intriguing framework for understanding the atypical PB2 sequence signatures of bat influenza viruses. The H17N10 polymerase contains serine substitutions at both the PB2-282^nd^ and 627^th^ residues that independently influence symmetric and asymmetric dimer formation, respectively (Fig. 5). Whereas the PB2-K627S substitution compromises interaction with human ANP32A and thereby affects asymmetric dimer formation, PB2-E282S weakens the Mid-NLS interaction and consequently impacts symmetric dimer formation. The two substitutions may therefore have evolved epistatically within bat virus polymerase to reshape the equilibrium between symmetric and asymmetric polymerase populations.

Such compensatory tuning of the polymerase dimeric states is consistent with previous observations from influenza viruses adapting to ANP32A/B-deficient cells^48^. In such a scenario, FluPol adapts to utilize the suboptimal ANP32E protein, resulting in a relatively destabilized asymmetric dimer, while simultaneously acquiring compensatory mutations that further destabilize the symmetric dimer interface. These observations, together with our findings, suggest that productive FluPol function depends not simply on formation of either dimeric state, but also on maintaining an appropriate equilibrium between symmetric and asymmetric dimer populations. The PB2-E282S substitution in bat influenza polymerase therefore represents a potential epistatic adaptation that couples conformational stability to the regulation of dimeric states, contributing to the restricted replication fitness of bat FluPol in non-bat host cells.

### Phosphomimetic adaptations that conferred fitness to the FluPol

The atypical serine residues at PB2-282 and 627 provide an additional layer of regulation linking polymerase sequence variation to host-dependent fitness. Recombinant H1N1 viruses carrying serine substitutions at the corresponding positions of PB2 exhibit impaired replication, consistent with disruption of the transcription–replication balance (Figs. 1h, 3 d-h). Intriguingly, both residues can undergo phosphorylation, justifying strong natural preference of FluPol for harboring negatively charged glutamic or aspartic acid at these positions of the PB2 (Figs. 1i, Supplementary Fig. 3). Importantly, the functional consequences of these substitutions are not equivalent. Phosphorylation at PB2-627 is supposed to be restricting because it disrupts the requirement of a positively charged lysine at this position for human ANP32-dependent asymmetric dimer formation^58,37^. In contrast, phosphorylation at PB2-282 should be rewarding as it could potentially compensate for the loss of the negative charge at this position and partially restore the Mid-NLS interaction. Consistent with this distinction, the PB2-E282S virus exhibits less replication defect than the PB2-K627S virus (Fig. 1h).

Whether the H17N10 polymerase undergoes regulated phosphorylation at these positions during infection of its natural bat host remains an important question. One possibility is that bat influenza viruses represent a divergent evolutionary solution in which serine residues at critical polymerase adaptation sites create a dependence on host kinase activity. In this model, phosphorylation would act as a reversible molecular switch that modulates the transition between transcription- and replication-competent states. In other influenza lineages, evolutionary acquisition of glutamic acid (Flu A, B, C) or aspartic acid (FluD) at these positions may have provided a phosphomimetic solution, allowing the polymerase to encode the negative charge constitutively and thereby reduce its dependence on host kinase regulation. Our findings pave the path towards a molecular-level understanding of the FluPol transition between transcription and replication states. The PB2 Mid-NLS interaction provides an intrinsic structural cue for stabilizing the replication conformation, while concentration-dependent oligomerization provides an additional regulatory layer, reinforcing this state through dimer formation. Together with phosphorylation, these mechanisms suggest that FluPol does not simply alternate between two predefined conformations; rather, it navigates a dynamic conformational landscape in which intramolecular contacts, oligomeric state, host-factor engagement, and post-translational modification are integrated to determine polymerase activity. Understanding these molecular cues would provide a basis for the rational designing of broad-spectrum polymerase inhibitors as next generation anti-influenza drugs.

## Materials and methods

### Cells and viruses

HEK293T, MDCK and HeLa cells were maintained in DMEM supplemented with 10% fetal bovine serum (FBS), 1× Pen-Strep and 1× GlutaMAX (Gibco). A549 cells were maintained in DMEM/F-12 supplemented with the same additives. All cells were cultured at 37°C in 5% CO₂.

Influenza A virus (IAV) A/WSN/1933 (H1N1), including a strain encoding C-terminally FLAG-tagged PB2 (WSN PB2-FLAG), and influenza B virus B/Brisbane/60/2008 were used for infections as indicated. Recombinant WSN viruses carrying PB2-E282S, PB2-K627S substitutions were generated as described earlier^39^.

### Plasmids

Viral genes were derived from A/WSN/1933 (H1N1), A/H17N10/little yellow-shouldered bat/Guatemala/164/2009 or B/Brisbane/60/2008. For H1N1 and influenza B, polymerase subunits were expressed using pcDNA3-based PB2-3×FLAG, PB1 and PA constructs; NP was expressed from pCDNA6.2-NP-V5 (H1N1) or pcDNA3.1-NP (influenza B). H17N10 polymerase and NP were expressed using pCMV-PB2-HA, pcDNA-PA-FLAG, pcDNA3.2-PB1-V5 and pcDNA3.2-NP-V5 constructs, kindly provided by A. Mehle^59^.

Polymerase activity was measured using pHH21-vNA-Luc and pHH21-cNA-Luc reporter plasmids containing the luciferase gene in negative-sense (vRNA) or positive-sense (cRNA) orientation, respectively, flanked by the 5′ and 3′ untranslated regions (UTRs) of the H1N1 NA segment and driven by the human RNA polymerase I promoter and terminator. Equivalent reporter systems were used for influenza B and H17N10 polymerases^60,39^.

For NanoBiT® complementation assays (Promega), H1N1 and H17N10 PB1 proteins were fused at their C-termini to LgBiT or SmBiT, while human ANP32A was fused to C-terminal SmBiT. Fusion constructs incorporated a 15-residue glycine–serine linker and were cloned into pBiT1.1-C [TK/LgBiT] or pBiT1.1-C [TK/SmBiT] vectors to assess polymerase dimerization and polymerase–ANP32A interactions.

### Site-directed mutagenesis

Mutations in H1N1 PB1 and PA and H17N10 PB2 were introduced using the QuikChange II Site-Directed Mutagenesis Kit (Agilent Technologies) with primers designed using QuikChange Primer Design software, according to the manufacturer’s instructions. All mutations were verified by Sanger sequencing.

### Polymerase activity assays

HEK293T cells seeded in 24-well plates were transfected in triplicate with plasmids expressing PB1, PB2-FLAG/HA, PA and NP-V5, together with the vNA-Luc or cNA-Luc reporter, using Lipofectamine 3000. At 18–24 h post-transfection, luciferase activity was measured using the Luciferase Assay System (Promega, E1500) and a GloMax 20/20 luminometer (Promega), according to the manufacturer’s instructions. Polymerase activity was expressed as a percentage of the WT PB2 control. Samples lacking PB2 expression served as negative controls.

### Primer-extension assays

Viral RNPs were reconstituted by co-transfecting plasmids expressing PB1, PB2, PA and NP together with a WSN-NA segment vRNA expression plasmid. Alternatively, mini-vRNPs were reconstituted using the NP77 template, comprising 77 nucleotides derived from the WSN NP-segment UTRs, and the polymerase subunits. At 36 h post-transfection, total RNA was extracted using TRIzol (Invitrogen). Viral mRNA, cRNA and vRNA were quantified by primer extension as previously described^39^, using [γ-^32^P]ATP-labelled strand-specific primers and a 5s rRNA-specific primer as an internal loading control (primer sequences in Supplementary Table 1). Extension products were resolved by 6% denaturing PAGE containing 7 M urea in 0.5X TBE and visualized using a Typhoon FLA-7000 phosphorimager. Band intensities were quantified using Image Lab, normalized to 5S rRNA, and expressed relative to the WT PB2 control using GraphPad Prism 9. The mRNA-to-vRNA ratio was calculated to assess the relative balance between transcription and replication.

### Multicycle replication kinetics

A549 cells at 80-90% confluency were infected in triplicates with wild-type or mutant Influenza A/H1N1/WSN/1933 (H1N1) viruses at a multiplicity of infection (M.O.I) of 0.001 in serum-free virus growth medium (VGM: 1X DMEM, 1X PenStrep, 4 mM GlutaMAX, 0.2% bovine serum albumin [BSA], 25 mM HEPES). Following adsorption, the inoculum was removed and pre-warmed VGM supplemented with 0.5 μg/mL TPCK-treated trypsin was added and incubated at 37°C in 5% CO_2_. Subsequently, culture supernatants were harvested at 8, 16, 24, 48 hours post infection and viral titers were determined by plaque assay on MDCK cells.

### LC-MS/MS analysis

A549 cells were infected with wild-type or mutant Influenza A/H1N1/WSN/1933 viruses expressing C-terminally FLAG-tagged PB2 (WSN-PB2-FLAG) at a M.O.I of 1. At 24 h post-infection, cells were lysed in Co-IP buffer (50 mM Tris-HCl, pH 7.4, 150 mM NaCl, and 0.5% NP-40) supplemented with protease and phosphatase inhibitors. RdRp complexes were affinity-purified using anti-FLAG M2 resin (Sigma, M2220), eluted in Laemmli sample buffer, and resolved by SDS-PAGE. Following Coomassie staining, protein bands at approximately 75 kDa were excised, destained, and subjected to in-gel trypsin digestion.

The resulting peptides were analysed using a Thermo Q Exactive HF-X mass spectrometer coupled to an Easy-nLC 1200 system. Peptides were separated at 300 nL/min on C18 trap (75 µm × 2 cm) and analytical (75 µm × 25 cm) columns using a 120-min gradient. MS data were acquired at 60,000 resolution (AGC target, 3 × 10⁶; maximum injection time, 50 ms; scan range, 400–1600 m/z). MS/MS spectra were acquired at 15,000 resolution (AGC target, 1 × 10⁵; maximum injection time, 50 ms) using data-dependent acquisition of the top 30 precursors, a 1.6 m/z isolation window, and a 25-s dynamic exclusion. The nanospray voltage was 1.8 kV, the capillary temperature was 250°C, and the funnel RF level was 40. The ion series for the phosphor-peptides of interest are provided in Supplementary Tables 2 and 3.

### Virus replication quantitation by sense-specific qRT-PCR

A549 cells at 80–90% confluency was synchronously infected with wild-type or mutant A/H1N1/WSN/1933 viruses at an M.O.I of 1. Cells were harvested at the indicated times post-infection, and total RNA was extracted using TRIzol reagent (Invitrogen). For quantification of viral v-, c- and mRNAs, we adapted a strand-specific RT-qPCR protocol described earlier ^61,62,44^, using primers listed in Supplementary Table 4. For each sample, 2 µg of RNA was reverse-transcribed using M-MuLV reverse transcriptase with strand-specific primers for NP-segment vRNA or cRNA, oligo(dT) for viral mRNA, or cellular 18s rRNA in separate reactions.

qPCR was performed in triplicate using iTaq Universal SYBR Green Supermix (Bio-Rad) on QuantStudio 5 Real-Time PCR System (Applied Biosystems). Viral RNA levels were normalized to cellular 18s rRNA using the ΔCt method and expressed relative to the corresponding 0-h post-infection levels using the ΔΔCt method. Mutant viral RNA levels were further expressed as percentages of wild-type levels at each time point. Data were analysed and plotted using GraphPad Prism 9.

### Co-immunoprecipitation assay

For polymerase dimerization assays, HEK293T cells were co-transfected with PB1, PA, and FLAG- and HA-tagged wild-type or mutant PB2. For polymerase–ANP32A interaction assays, cells were co-transfected with PB1, PA, PB2-FLAG (wild-type or mutant), and human ANP32A-V5. At 36 h post-transfection, cells were lysed in co-IP buffer (50 mM Tris-HCl, pH 7.4, 150 mM NaCl, and 0.5% NP-40) supplemented with protease and phosphatase inhibitor cocktails. Clarified lysates were pre-cleared with protein A agarose beads (Bio Bharati Life Science) and subjected to immunoprecipitation. Immune complexes were captured using protein A magnetic beads (Invitrogen Dynabeads), washed, and eluted in 1× Laemmli buffer before SDS-PAGE and western blotting. Co-immunoprecipitated protein levels were normalized to the corresponding immunoprecipitated protein by densitometry using Bio-Rad Image Lab and expressed as a percentage of the wild-type PB2 value using GraphPad Prism 9.

### cRNA stabilization assay

cRNA stabilization was assessed as previously described^21,63^. Briefly, HEK293T cells were transfected to pre-express the viral RdRP-PB2, PA, and wildtype or RNA-synthesis deficient mutant PB1 (445/446A), together with NP-v5. At 24 hours post transfection, cells were infected with Influenza A/H1N1/WSN/1933 (H1N1) virus at M.O.I. of 10 in presence of 1mM cycloheximide (CHX). Cells were harvested 6 hours post infection, total RNA was isolated using TRIzol and NA-segment based primer extension performed as described above to visualize and quantify viral mRNA, vRNA, and cRNA synthesis.

### *In vitro* ApG-primed replication initiation

The 77-nt NP-segment panhandle RNA (NP77), comprising the 45-nt 3′ UTR and 23-nt 5′ UTR of IAV/H1N1/WSN, was generated by *in vitro* transcription of annealed oligonucleotide-derived dsDNA using T7 RNA polymerase (Roche). vRNA- and cRNA-sense templates were transcribed using the appropriate T7 promoter orientation, and products were purified using TRIzol (primer sequences in Supplementary Table 5).

HEK293T cells transiently expressing wildtype or mutant PB2-FLAG, PB1, PA were lysed 36 hours post transfection in RNAse-free Co-IP buffer supplemented with 1X protease inhibitor cocktail and phosphatase inhibitor. RdRp complexes were immunoprecipitated using anti-FLAG magnetic beads, washed with RNase-free transcription buffer (10 mM Tris [pH 7.5], 75 mM NaCl, 5% glycerol, and 0.05% NP-40), and resuspended in 5 μl of the same buffer.

ApG-primed extension assays were performed as previously described ^43,47^. Bead-bound RdRP complexes were incubated for 1.5 h at 30°C in 20 μl reaction mixture containing 0.5 mM ApG (Jena Bioscience), 1 mM ATP, 0.5 mM GTP, 0.5 mM CTP, 0.5 μM cold UTP, 5 mM MgCl_2_, 1 mM DTT, 1 U/ μl RiboLock RNase Inhibitor (Thermo Fisher), 0.05 μM [α-^32^P] UTP and 1.5 μM NP77 template (vRNA- or cRNA-sense) in 1X transcription buffer with intermittent mixing. Reaction was terminated with 20 μl 2X RNA Loading Dye (95% formamide, 1 mM EDTA, 0.02% bromophenol blue and xylene cyanol) and resolved by 20% PAGE containing 7 M urea in 1X TBE, visualized by phosphorimaging and quantified by densitometric analysis, expressed as percentage wildtype.

### Split Nano-Luciferase assay

Split luciferase assays were adapted from Sheppard et al. to the Nano-luciferase based Promega NanoBiT technology^48^.To assess viral RdRP dimerization, HEK293T cells seeded in 48-well plates were transfected in triplicate with expression plasmids encoding H1N1 or H17N10 PB1 fused C-terminus Large BiT (LgBiT; 17.6kDa) and Small BiT (SmBiT; 11 amino acids) (0.04 μg each), together with PA (0.08 μg) and PB2 (0.08 μg). To assess RdRp-human ANP32A interactions, cells were transfected with plasmids encoding PB1-LgBiT, PB2, PA, and human ANP32A fused to C-terminal SmBiT (huANP32A-SmBiT), with 0.06 μg of each plasmid. Cells expressing PB1-LgBiT, PB1-SmBiT, or huANP32A-SmBiT in the absence of the corresponding complementary fusion partner were included as negative controls. At 24 hours post-transfection, cells were harvested and luminescence was measured using Nano-Glo Luciferase Assay System (Promega) according to manufacturer’s instructions. Luminescence signals obtained with mutant PB2 proteins were normalized to wild-type PB2 protein and expressed as a percentage.

### Immunofluorescence and quantification of PB2 nuclear localization

HeLa cells grown on coverslips were transfected with plasmids encoding wildtype or mutant PB2-FLAG using Lipofectamine 3000. At 24 hours post-transfection, monolayers were washed with PBS and fixed with 3% formaldehyde. The fixative was quenched with 0.1 M glycine and cells were permeabilized with 0.1% Triton-X 100, followed by blocking with 3% BSA. Cells were then incubated overnight at 4℃ with anti-FLAG primary antibody (1:1000) diluted in 3% BSA in PBS. PB2-FLAG was detected using Alexa Fluor-488 conjugated anti-mouse secondary antibody (1:1000; Invitrogen) and nuclei were counterstained with DAPI (1 μg/mL). Images were acquired using 60X oil-immersion objective on an Olympus FV3000 confocal microscope with 488-nm laser excitation and processed using ImageJ. At least 50 cells (n = 50) were randomly selected from 3 different fields for each experimental set. Data are presented as mean ± standard deviation (n = 50) and plotted using GraphPad Prism 9.

### Phylogenetic analysis and multiple sequence alignments (MSA)

Complete protein sequences of influenza A PB1, PB2, PA, grouped by host organism, were retrieved from NCBI Influenza Virus Database (https://www.ncbi.nlm.nih.gov/labs/virus/vssi/#/) and clustered at 85% identity using MMseqs2 easy-linclust tool available through Galaxy platform^64^. Relevant vaccine strains, lineage defining, laboratory and pandemic strains were manually added to generate a final dataset of ∼250 sequences per viral protein across diverse host range. Sequences were subjected to multiple sequence alignments (MSA) using Clustal Omega^65^, unrooted maximum likelihood phylogenetic trees were constructed using IQ-Tree and colour-coded representations were created using Rainbow Tree (https://www.hiv.lanl.gov/content/sequence/RAINBOWTREE/rainbowtree.html).

For extended alignment of PB2 Mid-domain and NLS, PB2 sequences from influenza A, B, C, D viruses were retrieved from NCBI Influenza Virus Database and consensus sequence generated for each virus type using Clustal Omega and were aligned using ESPrit3.2^66^.

### Structural analysis

Available PDB structures of influenza polymerase complex were analysed using PyMol (version 3.0; Schrödinger, LLC) or UCSF ChimeraX (version 1.11.1;^67^. Surface charge representations were generated using the APBS electrostatics module integrated with PyMol^68^. In silico mutagenesis, interatomic distance measurements and structural alignments were performed using UCSF ChimeraX.

### Statistical analysis

All experiments were performed in triplicate, with each experiment independently repeated at least three times. Data were analysed as indicated and plotted using GraphPad Prism 9, expressed as mean ± standard deviation (SD; n = 3). Statistical comparisons were performed using two-tailed Student’s t-test, or one-way or two-way analysis of variance (ANOVA), as appropriate for the experimental design. A *P*-value < 0.001 was considered statistically significant. Exact *P*-values are indicated above the corresponding bars in the graphs.

## Acknowledgements

We thank Prof. Andrew Mehle (University of Wisconsin Madison) for providing important plasmids and for his valuable insights on the manuscript. The project is not supported by any funding. However, A.M. acknowledges financial support to the lab from Anusandhan National Research Foundation (ANRF)(SERB-DST (CRG/2022/003628), Department of Biotechnology (DBT)(BT-BT/PR58658/BMS2/156/208/2025 and BT/PR65384/BIOAH/164/172/2026), Indian Council of Medical Research (ICMR) (EMDR/SG/14/2025-00024) and West Bengal Department of Science and Technology (WBDST) (1111(Sanc.)/STBT-11012(12)/2/2024-WBSCST SEC). A.D. acknowledges support from the Prime Minister’s Research Fellowship (PMRF), awarded by the Ministry of Education, Government of India. We thank Soumik Dey for his assistance with the immunofluorescence studies. We also thank Ms. Elizabeth Remily-Wood, Director of the USF MCOM Proteomics Facility. The proteomics work was supported, in part, by the Morsani College of Medicine (MCOM), University of South Florida (USF).

## Author Contributions

A.D. and A. M. conceived and designed the study. A.D. performed the experiments and analyzed the data, with inputs from A.M. A.R. performed the mass spectrometry experiments and was responsible for data acquisition, processing and analysis. A.M. supervised the project. A.D. and A.M. wrote the manuscript with input from all authors. All authors reviewed and approved the final version of the manuscript.

## Data Availability

Whole-genome sequencing of the wild-type A/India/H1N1-P2/2020 (H1N1) and mutant A/India/H1N1-P3/2020 (H1N1; PB2-E282S) viruses was performed by next-generation sequencing (NGS), as described previously.^39^

The corresponding nucleotide sequences have been deposited in GenBank under the accession numbers listed below.

| Virus | Mutations | Protein | GenBank ID |
|---|---|---|---|
| A/India/H1N1-P2/2020(H1N1) | None | PB2, PB1, PA, HA, NP, NA, M, NS | OL831179, OL831180, OL831181, OL831182, OL831183, OL831184, OL831185, OL831186 |
| A/India/H1N1-P3/2020(H1N1) | PB2-E282S | PB2, PB1, PA, HA, NP, NA, M, NS | OL831157, OL831158, OL831159, OL831160, OL831161, OL831162, OL831163, OL831164 |

For the PB2-K627S/WSN mutant virus sanger sequencing was performed to confirm the sequence of all three polymerase subunits (PB1, PB2 and PA) showing only intended mutation in PB2 ORF.

## References

1. Taubenberger, J. K. & Kash, J. C. Influenza Virus Evolution, Host Adaptation, and Pandemic Formation. Cell Host Microbe 7, 440–451 (2010).

2. Mostafa, A., Abdelwhab, E. M., Mettenleiter, T. C. & Pleschka, S. Zoonotic Potential of Influenza A Viruses: A Comprehensive Overview. Viruses 10, 497 (2018).

3. Chrzastek, K., Lieber, C. M. & Plemper, R. K. H5N1 Clade 2.3.4.4b: Evolution, Global Spread, and Host Range Expansion. Pathogens 14, 929 (2025).

4. Mifsud, E. J., Hayden, F. G. & Hurt, A. C. Antivirals targeting the polymerase complex of influenza viruses. Antiviral Res. 169, 104545 (2019).

5. Subbarao, E. K., London, W. & Murphy, B. R. A single amino acid in the PB2 gene of influenza A virus is a determinant of host range. J. Virol. 67, 1761–1764 (1993).

6. Mehle, A. & Doudna, J. A. Adaptive strategies of the influenza virus polymerase for replication in humans. Proc. Natl. Acad. Sci. 106, 21312–21316 (2009).

7. Zhu, W. et al. Mutations in Polymerase Genes Enhanced the Virulence of 2009 Pandemic H1N1 Influenza Virus in Mice. PLOS ONE 7, e33383 (2012).

8. Chin, A. W. H., Leong, N. K. C., Nicholls, J. M. & Poon, L. L. M. Characterization of influenza A viruses with polymorphism in PB2 residues 701 and 702. Sci. Rep. 7, 11361 (2017).

9. Arranz, R. et al. The Structure of Native Influenza Virion Ribonucleoproteins. Science 338, 1634–1637 (2012).

10. Moeller, A., Kirchdoerfer, R. N., Potter, C. S., Carragher, B. & Wilson, I. A. Organization of the Influenza Virus Replication Machinery. Science 338, 1631–1634 (2012).

11. Peng, R. et al. Molecular basis of influenza ribonucleoprotein complex assembly and processive RNA synthesis. Science 388, eadq7597 (2025).

12. Dou, D., Revol, R., Östbye, H., Wang, H. & Daniels, R. Influenza A Virus Cell Entry, Replication, Virion Assembly and Movement. Front. Immunol. 9, (2018).

13. Fodor, E. The RNA polymerase of influenza a virus: mechanisms of viral transcription and replication. Acta Virol. 57, 113–122 (2013).

14. Reich, S. et al. Structural insight into cap-snatching and RNA synthesis by influenza polymerase. Nature 516, 361–366 (2014).

15. Guilligay, D. et al. The structural basis for cap binding by influenza virus polymerase subunit PB2. Nat. Struct. Mol. Biol. 15, 500–506 (2008).

16. Yuan, P. et al. Crystal structure of an avian influenza polymerase PAN reveals an endonuclease active site. Nature 458, 909–913 (2009).

17. Fodor, E. & te Velthuis, A. J. W. Structure and Function of the Influenza Virus Transcription and Replication Machinery. Cold Spring Harb. Perspect. Med. 10, a038398 (2020).

18. Zhu, Z., Fodor, E. & Keown, J. R. A structural understanding of influenza virus genome replication. Trends Microbiol. 31, 308–319 (2023).

19. Deng, T., Vreede, F. T. & Brownlee, G. G. Different de novo initiation strategies are used by influenza virus RNA polymerase on its cRNA and viral RNA promoters during viral RNA replication. J. Virol. 80, 2337–2348 (2006).

20. Oymans, J. & Te Velthuis, A. J. W. A Mechanism for Priming and Realignment during Influenza A Virus Replication. J. Virol. 92, e01773–17 (2018).

21. Vreede, F. T., Jung, T. E. & Brownlee, G. G. Model suggesting that replication of influenza virus is regulated by stabilization of replicative intermediates. J. Virol. 78, 9568–9572 (2004).

22. Jorba, N., Coloma, R. & Ortín, J. Genetic trans-Complementation Establishes a New Model for Influenza Virus RNA Transcription and Replication. PLOS Pathog. 5, e1000462 (2009).

23. York, A., Hengrung, N., Vreede, F. T., Huiskonen, J. T. & Fodor, E. Isolation and characterization of the positive-sense replicative intermediate of a negative-strand RNA virus. Proc. Natl. Acad. Sci. 110, E4238–E4245 (2013).

24. Thierry, E. et al. Influenza Polymerase Can Adopt an Alternative Configuration Involving a Radical Repacking of PB2 Domains. Mol. Cell 61, 125–137 (2016).

25. Wandzik, J. M., Kouba, T. & Cusack, S. Structure and Function of Influenza Polymerase. Cold Spring Harb. Perspect. Med. 11, a038372 (2021).

26. Pflug, A., Lukarska, M., Resa-Infante, P., Reich, S. & Cusack, S. Structural insights into RNA synthesis by the influenza virus transcription-replication machine. Virus Res. 234, 103– 117 (2017).

27. Tarendeau, F. et al. Structure and nuclear import function of the C-terminal domain of influenza virus polymerase PB2 subunit. Nat. Struct. Mol. Biol. 14, 229–233 (2007).

28. Tarendeau, F. et al. Host Determinant Residue Lysine 627 Lies on the Surface of a Discrete, Folded Domain of Influenza Virus Polymerase PB2 Subunit. PLOS Pathog. 4, e1000136 (2008).

29. Dias, A. et al. The cap-snatching endonuclease of influenza virus polymerase resides in the PA subunit. Nature 458, 914–918 (2009).

30. Pflug, A., Guilligay, D., Reich, S. & Cusack, S. Structure of influenza A polymerase bound to the viral RNA promoter. Nature 516, 355–360 (2014).

31. Kouba, T., Drncová, P. & Cusack, S. Structural snapshots of actively transcribing influenza polymerase. Nat. Struct. Mol. Biol. 26, 460–470 (2019).

32. Delaforge, E. et al. Large-Scale Conformational Dynamics Control H5N1 Influenza Polymerase PB2 Binding to Importin α. J. Am. Chem. Soc. 137, 15122–15134 (2015).

33. Li, H. et al. An intermediate state allows influenza polymerase to switch smoothly between transcription and replication cycles. Nat. Struct. Mol. Biol. 30, 1183–1192 (2023).

34. Fan, H. et al. Structures of influenza A virus RNA polymerase offer insight into viral genome replication. Nature 573, 287–290 (2019).

35. Carrique, L. et al. Host ANP32A mediates the assembly of the influenza virus replicase. Nature 587, 638–643 (2020).

36. Staller, E. et al. Structures of H5N1 influenza polymerase with ANP32B reveal mechanisms of genome replication and host adaptation. Nat. Commun. 15, 4123 (2024).

37. Arragain, B. et al. Structures of influenza A and B replication complexes give insight into avian to human host adaptation and reveal a role of ANP32 as an electrostatic chaperone for the apo-polymerase. Nat. Commun. 15, 6910 (2024).

38. Wang, F. et al. The C-terminal LCAR of host ANP32 proteins interacts with the influenza A virus nucleoprotein to promote the replication of the viral RNA genome. Nucleic Acids Res. 50, 5713–5725 (2022).

39. Banerjee, S. et al. The Species-Specific 282 Residue in the PB2 Subunit of the Polymerase Regulates RNA Synthesis and Replication of Influenza A Viruses Infecting Bat and Nonbat Hosts. J. Virol. 96, e02190-21 (2022).

40. Tong, S. et al. A distinct lineage of influenza A virus from bats. Proc. Natl. Acad. Sci. 109, 4269–4274 (2012).

41. Tong, S. et al. New World Bats Harbor Diverse Influenza A Viruses. PLOS Pathog. 9, e1003657 (2013).

42. Ciminski, K., Pfaff, F., Beer, M. & Schwemmle, M. Bats reveal the true power of influenza A virus adaptability. PLOS Pathog. 16, e1008384 (2020).

43. te Velthuis, A. J. W., Long, J. S. & Barclay, W. S. Assays to Measure the Activity of Influenza Virus Polymerase. in Influenza Virus: Methods and Protocols (ed. Yamauchi, Y.) 343–374 (Springer, New York, NY, 2018). doi:10.1007/978-1-4939-8678-1_17.

44. Jana, I. D., Dey, S., Si, M., Roy, A. & Mondal, A. Host Protein Kinase C*a*: The novel Mitogen Activated Protein Kinase (MAPK) specific scaffold regulating nuclear export of influenza virus ribonucleoprotein complexes. PLOS Pathog. 21, e1013841 (2025).

45. Nilsson-Payant, B. E., tenOever, B. R. & te Velthuis, A. J. W. The Host Factor ANP32A Is Required for Influenza A Virus vRNA and cRNA Synthesis. J. Virol. 96, e02092–21 (2022).

46. Mänz, B., Brunotte, L., Reuther, P. & Schwemmle, M. Adaptive mutations in NEP compensate for defective H5N1 RNA replication in cultured human cells. Nat. Commun. 3, 802 (2012).

47. te Velthuis, A. J. W., Robb, N. C., Kapanidis, A. N. & Fodor, E. The role of the priming loop in influenza A virus RNA synthesis. Nat. Microbiol. 1, 16029 (2016).

48. Sheppard, C. M. et al. An Influenza A virus can evolve to use human ANP32E through altering polymerase dimerization. Nat. Commun. 14, 6135 (2023).

49. Long, J. S. et al. Species specific differences in use of ANP32 proteins by influenza A virus. eLife 8, e45066 (2019).

50. Baker, S. F., Ledwith, M. P. & Mehle, A. Differential Splicing of ANP32A in Birds Alters Its Ability to Stimulate RNA Synthesis by Restricted Influenza Polymerase. Cell Rep. 24, 2581–2588.e4 (2018).

51. Keown, J. R. et al. Mapping inhibitory sites on the RNA polymerase of the 1918 pandemic influenza virus using nanobodies. Nat. Commun. 13, 251 (2022).

52. Resa-Infante, P. et al. The Host-Dependent Interaction of α-Importins with Influenza PB2 Polymerase Subunit Is Required for Virus RNA Replication. PLOS ONE 3, e3904 (2008).

53. Lukarska, M. et al. Structural basis of an essential interaction between influenza polymerase and Pol II CTD. Nature 541, 117–121 (2017).

54. Krischuns, T. et al. The host RNA polymerase II C-terminal domain is the anchor for replication of the influenza virus genome. Nat. Commun. 15, 1064 (2024).

55. Seong, B. L., Kobayashi, M., Nagata, K., Brownlee, G. G. & Ishihama, A. Comparison of Two Reconstituted Systems for In Vitro Transcription and Replication of Influenza Virus. J. Biochem. (Tokyo) 111, 496–499 (1992).

56. Zhu, Z., Fan, H. & Fodor, E. Defining the minimal components of the influenza A virus replication machinery via an in vitro reconstitution system. PLOS Biol. 21, e3002370 (2023).

57. Chang, S. et al. Cryo-EM Structure of Influenza Virus RNA Polymerase Complex at 4.3 Å Resolution. Mol. Cell 57, 925–935 (2015).

58. Camacho-Zarco, A. R. et al. Molecular basis of host-adaptation interactions between influenza virus polymerase PB2 subunit and ANP32A. Nat. Commun. 11, 3656 (2020).

59. Poole, D. S. et al. Influenza A Virus Polymerase Is a Site for Adaptive Changes during Experimental Evolution in Bat Cells. J. Virol. 88, 12572–12585 (2014).

60. Kedia, N., Banerjee, S. & Mondal, A. A Comprehensive Roadmap Towards the Generation of an Influenza B Reporter Assay Using a Single DNA Polymerase-Based Cloning of the Reporter RNA Construct. Front. Microbiol. 13, (2022).

61. Kawakami, E. et al. Strand-specific real-time RT-PCR for distinguishing influenza vRNA, cRNA, and mRNA. J. Virol. Methods 173, 1–6 (2011).

62. Kuchipudi, S. V. et al. 18S rRNAis a reliable normalisation gene for real time PCR based on influenza virus infected cells. Virol. J. 9, 230 (2012).

63. Das, S. et al. Influenza virus Non Structural protein 1 (NS1) Chaperones Nucleoprotein (NP) Oligomerization to Coordinate Ribonucleoprotein Assembly and Genome Replication. 2026.05.16.725629 Preprint at 10.64898/2026.05.16.725629 (2026).

64. Steinegger, M. & Söding, J. Clustering huge protein sequence sets in linear time. Nat. Commun. 9, 2542 (2018).

65. Sievers, F. et al. Fast, scalable generation of high-quality protein multiple sequence alignments using Clustal Omega. Mol. Syst. Biol. 7, MSB201175 (2011).

66. Gouet, P., Courcelle, E., Stuart, D. I. & M√©toz, F. ESPript: analysis of multiple sequence alignments in PostScript. Bioinformatics 15, 305–308 (1999).

67. Meng, E. C. et al. UCSF ChimeraX: Tools for structure building and analysis. Protein Sci. Publ. Protein Soc. 32, e4792 (2023).

68. Jurrus, E. et al. Improvements to the APBS biomolecular solvation software suite. Protein Sci. Publ. Protein Soc. 27, 112–128 (2018).

