## Supplementary Figures 1-10 and Supplementary Tables 1-5 for "Decoding the molecular cues regulating the transition of influenza virus polymerase from transcriptase to replicase"

### Supplementary Figure 1

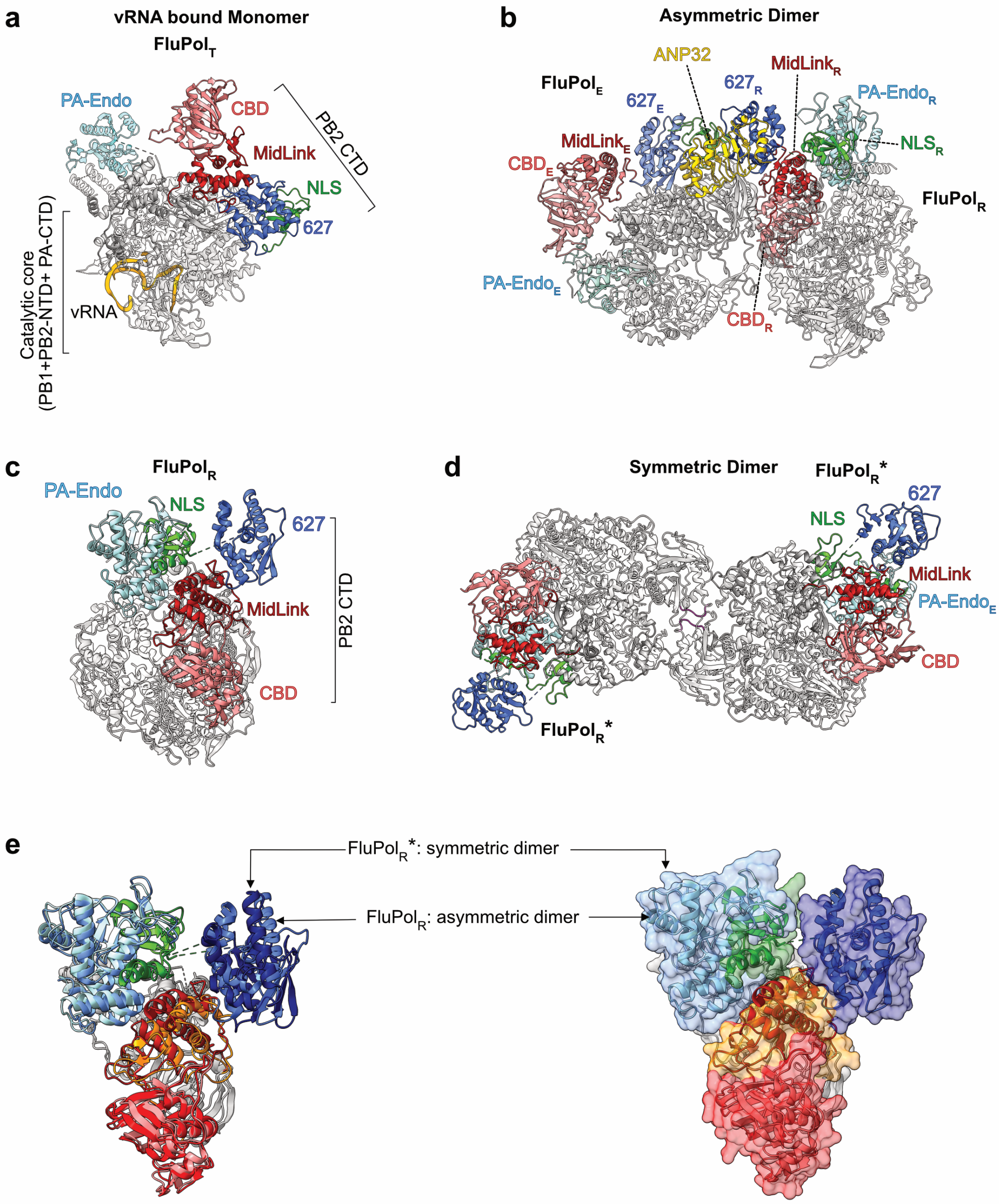


**Supplementary Figure 1**. (a-e) Structures of influenza A polymerase in various functional states representing (a) H17N10 polymerase in transcriptase mode (PDB ID 4WSB) (b) asymmetric dimer of H5N1 polymerase in complex with human ANP32B (PDB ID 8R1J) (c) FluPol$_R$ from the asymmetric-dimer complex (d) symmetric dimer of H5N1 polymerase (PDB ID 6QPF) showing the replicase-like organization of PB2-CTD (FluPol$_R$*) (e) structural alignment of PB2 from the asymmetric and symmetric dimers, both in cartoon representation (left) or showing FluPol$_R$ from the asymmetric-dimer in cartoon and FluPol$_R$* from the symmetric-dimer in surface representation (right). PB2 RMSD between 285 pruned atom pairs is 1.272 Å (across all 757 pairs; 5.564 Å).

**Supplementary Figure 2**

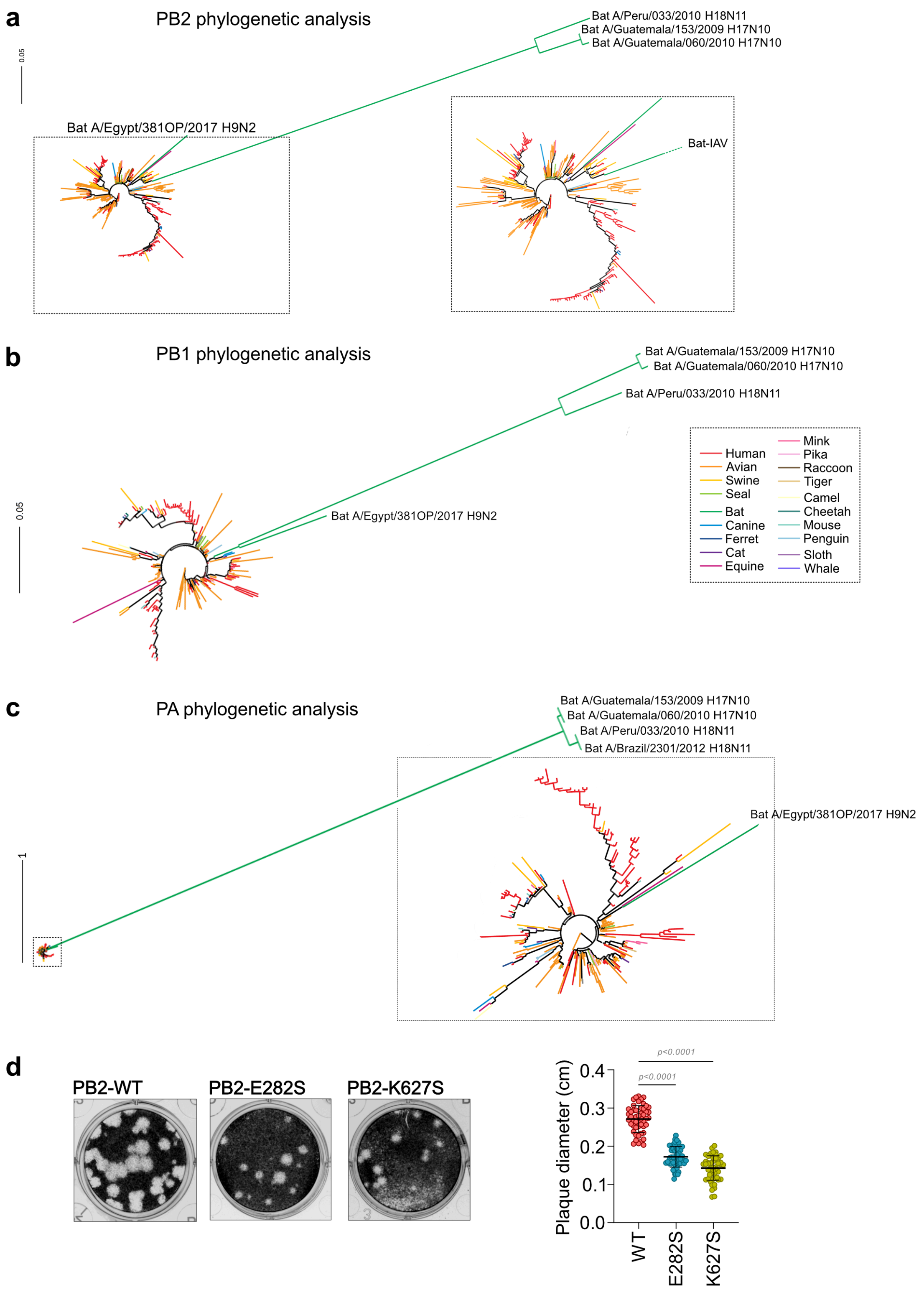


**Supplementary Figure 2.** (a-c) Phylogenetic analysis of PB2, PB1 and PA proteins displaying genetic divergence of the bat-IAV H17N10 and H18N11 polymerases. (d) Images of plaques obtained for WT or PB2-mutant A/H1N1/WSN/1933 viruses and quantification of their mean plaque diameters.

### Supplementary Figure 3

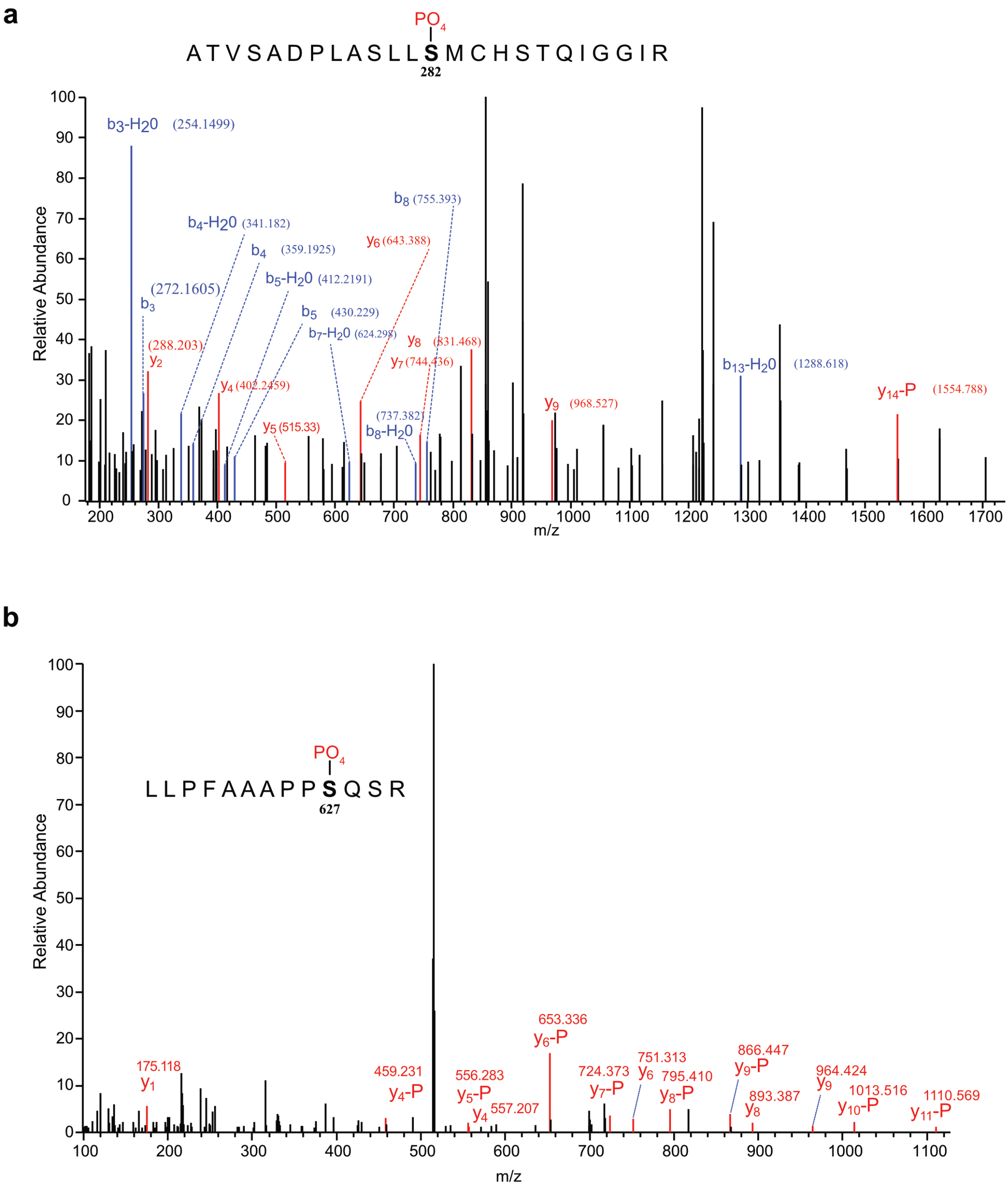


**Supplementary Figure 3.** LC-MS/MS analysis of the polymerase purified from A/H1N1/WSN PB2-E282S and PB2-K627S viruses. Chromatograms of phospho-peptides containing the phospho-serine at PB2-282nd (a) and PB2-627th (b) residues are displayed.

**Supplementary Figure 4**

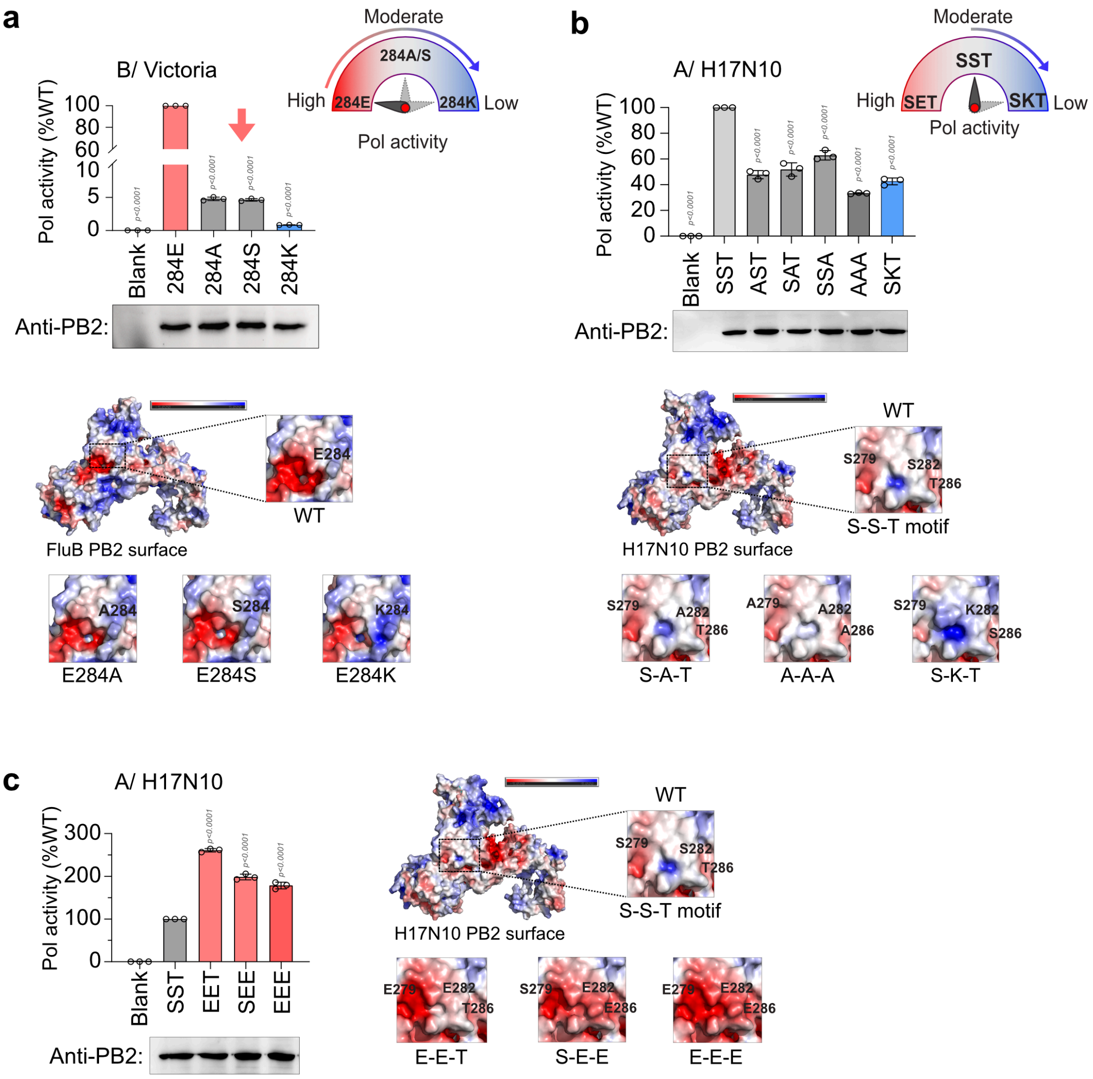


**Supplementary Figure 4.** (a) Polymerase activity of Influenza B/Brisbane bearing mutations at the PB2-E284 residue, normalized to WT. (b-c) Polymerase activity of A/H17N10 polymerase bearing alanine substitutions (b) or multiple glutamic acid (c) within the native PB2-"S279-S282-T286" motif. Schematics indicate the loss of polymerase activity; lower panels show the corresponding PB2 subunits in surface-charge representation; insets highlight mutation-induced changes in surface charge (PDB ID: 4WSB for A/H17N10, 4WSA for B/Brisbane).

**Supplementary Figure 5**

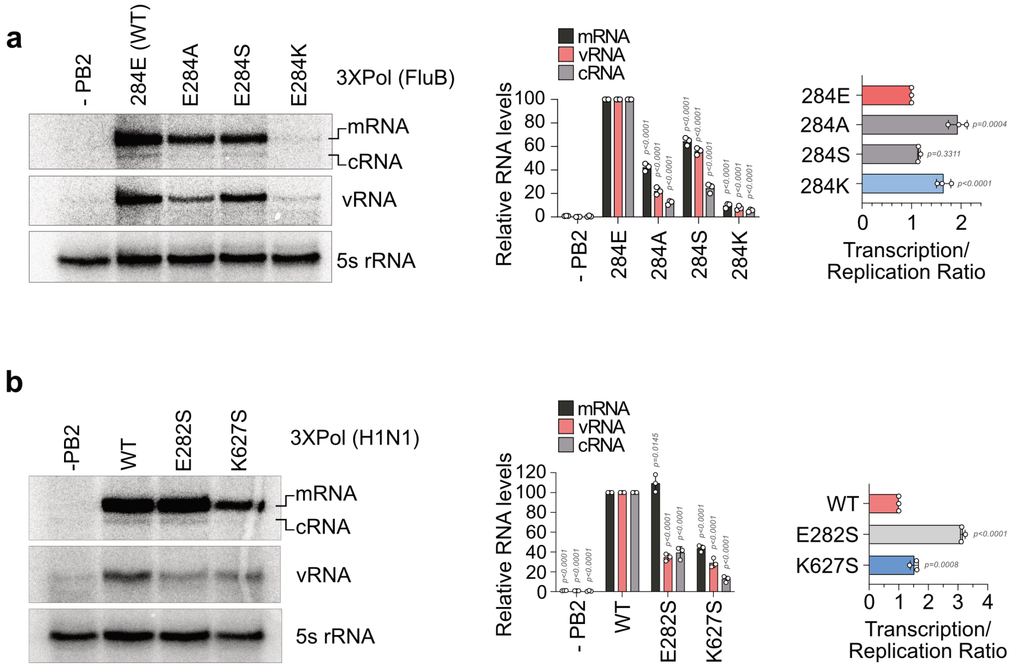


**Supplementary Figure 5.** Primer extension assays showing transcription and replication activities of (a) B/Brisbane harboring PB2-E284A, E282S, or E282K mutations; (b) H1N1 polymerase harboring PB2-E282S or PB2-K627S mutations, compared to the respective WT polymerase. Relative abundances of viral transcripts (mRNA) and replicons (c-/vRNAs) are shown, along with mRNA/vRNA ratio as a measure of the transcription–replication balance.

**Supplementary Figure 6**

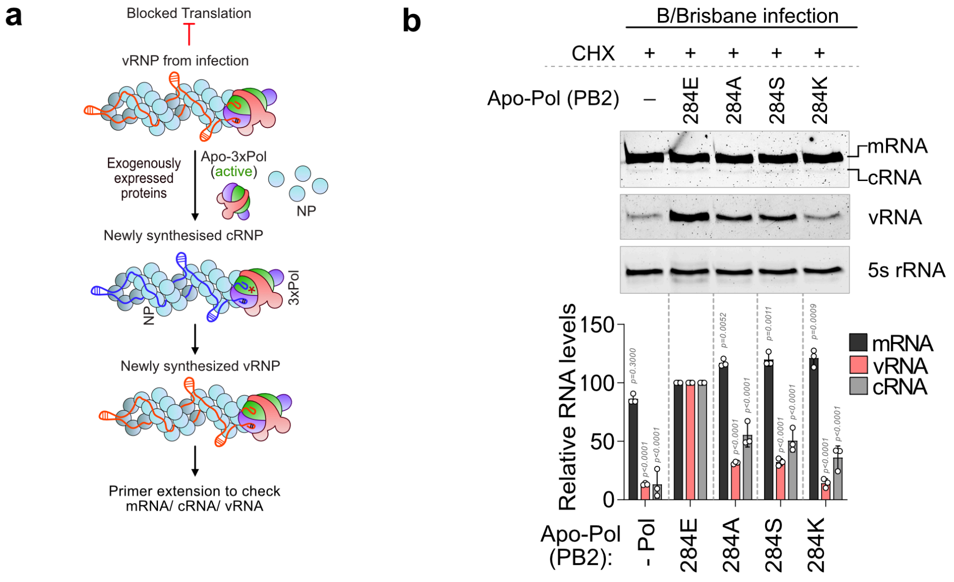


**Supplementary Figure 6.** Cells pre-expressing NP and polymerase harboring either WT or E284A/S/K-mutant PB2 were infected with B/Brisbane/60/2008 virus and treated with cycloheximide at 0 h.p.i. Viral RNA synthesis measured by primer extension.

**Supplementary Figure 7**

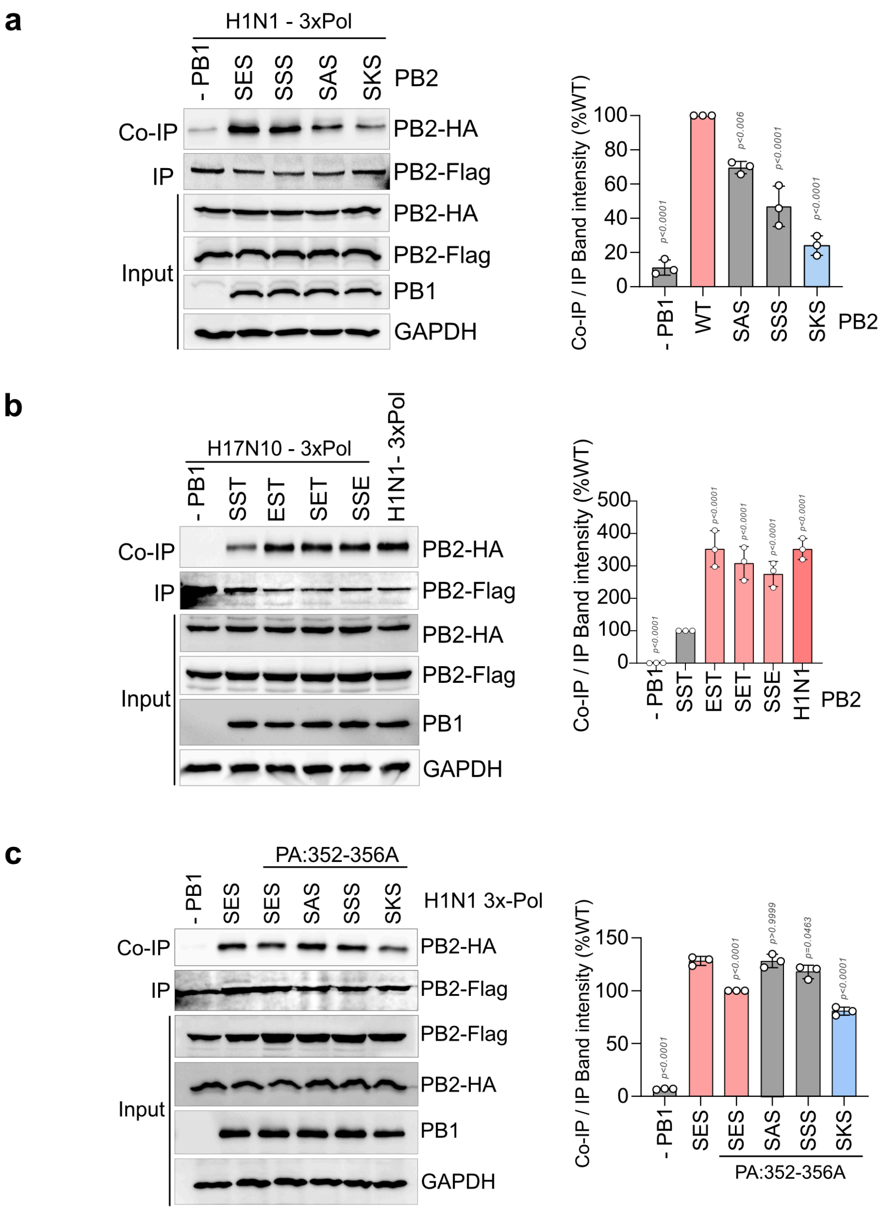


**Supplementary Figure 7.** FLAG or HA-epitope tagged PB2 (WT or mutant) along with PB1 and PA proteins were expressed in HEK293T cells. Polymerase dimerization was monitored by immunoprecipitating FLAG-tagged PB2 and estimating the co-precipitation of HA-tagged PB2. No PB1 set was included as negative control, as the efficient assembly of the heterotrimeric RdRp is a pre-requisite for the formation of dimer of heterotrimers. (a) Effect of PB2-“S-E-S” motif mutants were assessed upon dimerization ability of the H1N1 polymerase. (b) Effect of introduction of glutamic acid at different positions of the PB2-“S-S-S” motif were assessed upon H17N10 polymerase dimerization. (c) Same experiment as (a), except that the symmetric dimerization mutant PA-352-356A was used to assess the effect of PB2-“S-E-S” motif mutants on the asymmetric/ non-symmetric dimer formation ability of the polymerase.

**Supplementary Figure 8**

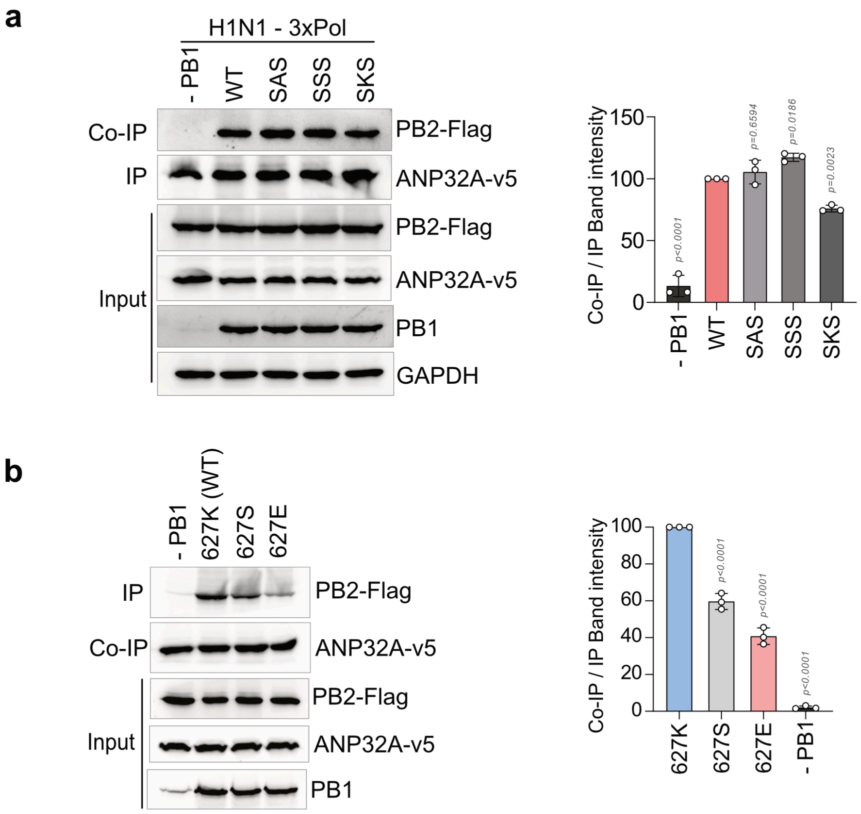


**Supplementary Figure 8.** V5-epitope tagged human ANP32A along with FLAG-epitope tagged PB2 (WT or mutant), untagged PB1 and PA proteins were expressed in HEK293T cells. Polymerase-ANP32 interaction was monitored by immunoprecipitating V5-tagged ANP32 and estimating the co-precipitation of FLAG-tagged PB2. (A) The effect of PB2-“S-E-S” motif mutants or (B) PB2-K627S or K627E mutants on polymerase-ANP32 interaction was assessed.

### Supplementary Figure 9

a

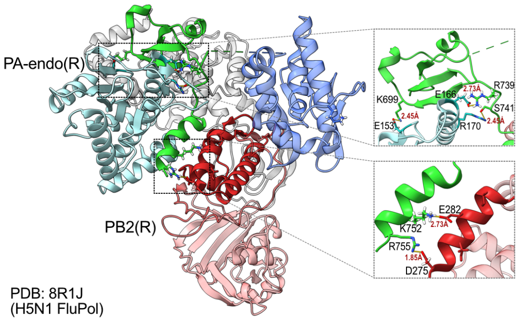


| PB2 (R)- Mid : NLS interactions | | |
|---|---|---|
| Res 1 (Mid) | Res 2 (NLS) | Distance (Å) |
| GLU 282 | LYS 752 | 2.729 |
| ASP 275 | ARG 755 | 1.851 |
| **PB2 (R)-NLS : PA-endo(R) interactions** | | |
| LYS 699 | GLU 153 | 2.454 |
| ARG 739 | GLU 166 | 2.733 |
| SER 741 | ARG 170 | 2.448 |

b

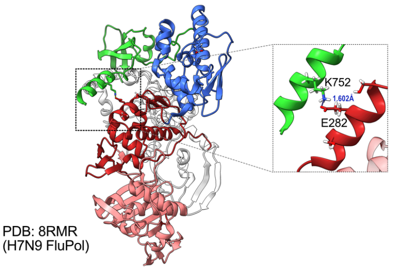


c

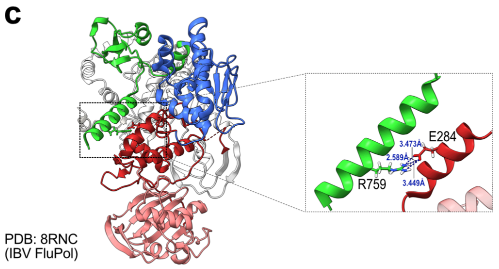


| FluPol | Res 1 (Mid) | Res 2 (NLS) | Distance (Å) | Predicted Bond |
|---|---|---|---|---|
| H7N9 | GLU 282 | LYS 752 | 1.602 | salt-bridge |
| Flu B | GLU 284 | ARG 759 | 2.5 - 3.5 | salt-bridge |

d

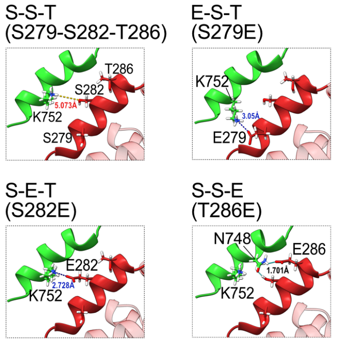


| Res 1 (Mid) | Res 2 (NLS) | Distance (Å) | Predicted Bond |
|---|---|---|---|
| SER 282 | LYS 752 | 5.073 | — |
| GLU 279 | LYS 752 | 3.050 | salt-bridge |
| GLU 282 | LYS 752 | 2.728 | salt-bridge |
| GLU 286 | ASN 748 | 1.701 | H-bond |

**Supplementary Figure 9.** (a) Structure of the PA-endonuclease domain and PB2 of IAV/H5N1 FluPol in replicase conformation ($FluPol_R$) highlighting potential conserved interactions (salt bridges or H-bonds) between PA-endo (light blue) and PB2-Mid domain (red), and between PB2-Mid and NLS-domains (green). (b-c) Structural analysis of potential interactions between the PB2-Mid (red) and NLS (green) domains specific to the replicase conformation ($FluPol_R$) in A/H7N9 (PDB ID 8RMR) (b) and influenza B (PDB ID 8RNC) (c) virus polymerases. Corresponding interatomic distances are provided in the accompanying tables. (d) Structural representation of bat-IAV "S279-S282-T286" (S-S-T) motif introduced into PB2 Mid-domain in the H5N1-$FluPol_R$ (PDB ID 8R1J), along with glutamic acid substitutions at each position and their potential interactions with PB2-K752/N748 in the NLS, as predicted using UCSF Chimera.

### Supplementary Figure 10

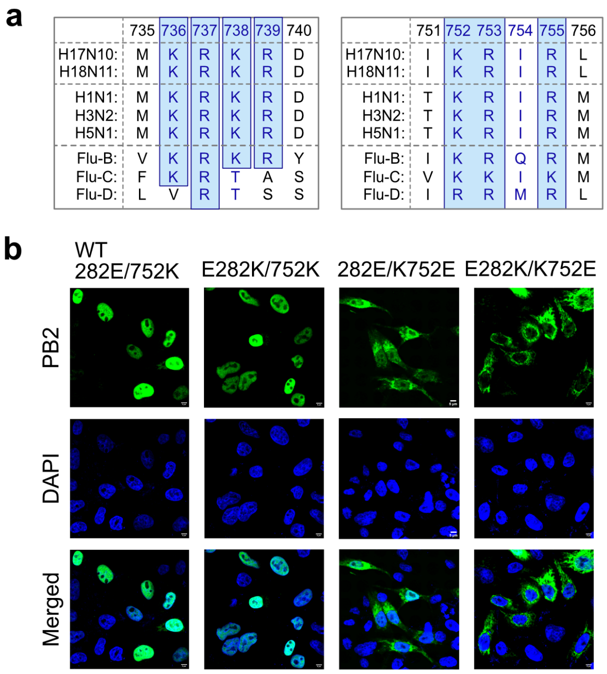


**Supplementary Figure 10**. (A) Multiple sequence alignment of PB2 NLS1 (residues 736-739) and NLS2 (residues 752-755) regions across influenza A, B, C, D viruses (B) Immunofluorescence analysis of nuclear localization of WT PB2 and mutants with substitutions at either 282, 752 or both positions.

**Supplementary Tables**

Supplementary Table 1. Sequences of primers used for primer extension:

| Primer name | Sequence (5’->3’) |
|---|---|
| NP77_vRNA | TGATTTCGATGTCACTCTGTGAG |
| Np77_mRNA | GCAGGGTAGATAATCACTCACAG |
| H1N1_NA_vRNA | TGGACTAGTGGGAGCATCAT |
| H1N1_NA_m/cRNA | TCCAGTATGGTTTTGATTTCCG |
| IBV_NA_vRNA | GTTGACATGGCTCTGTAATG |
| IBV_NA_m/cRNA | CCGAATATAGTAAGTATGATAATGAAG |
| 5s rRNA | TCCCAGGCGGTCTCCCATCC |

Supplementary Table 2. Ion series table for peptide ATVSADPLASLLsMCHSTQIGGIR:

| $b^+$ | $b\text{-}H_2O^+$ | Seq. | $y^+$ | $y^+$-P |
|---|---|---|---|---|
| 72.04439 | | A | | |
| 173.0921 | 155.0815 | T | 2494.183 | 2396.206 |
| 272.1605 | 254.1499 | V | 2393.135 | 2295.158 |
| 359.1925 | 341.182 | S | 2294.067 | 2196.09 |
| 430.2296 | 412.2191 | A | 2207.035 | 2109.058 |
| 545.2566 | 527.246 | D | 2135.998 | 2038.021 |
| 642.3093 | 624.2988 | P | 2020.971 | 1922.994 |
| 755.3934 | 737.3828 | L | 1923.918 | 1825.941 |
| 826.4305 | 808.4199 | A | 1810.834 | 1712.857 |
| 913.4625 | 895.452 | S | 1739.797 | 1641.82 |
| 1026.547 | 1008.536 | L | 1652.765 | 1554.788 |
| 1139.631 | 1121.62 | L | 1539.681 | 1441.704 |
| 1306.629 | 1288.618 | S-Phospho | 1426.597 | 1328.62 |
| 1437.67 | 1419.659 | M | 1259.598 | |
| 1597.7 | 1579.69 | C-Carbamidomethyl | 1128.558 | |
| 1734.759 | 1716.749 | H | 968.5272 | |
| 1821.791 | 1803.781 | S | 831.4683 | |
| 1922.839 | 1904.828 | T | 744.4363 | |
| 2050.897 | 2032.887 | Q | 643.3886 | |
| 2163.981 | 2145.971 | I | 515.33 | |
| 2221.003 | 2202.992 | G | 402.2459 | |
| 2278.024 | 2260.014 | G | 345.2245 | |
| 2391.108 | 2373.098 | I | 288.203 | |
| | | R | 175.119 | |

Supplementary Table 3. Ion series table for peptide LLPFAAAPPSQSR:

| $b^+$ | Seq. | $y^+$ | $y^+$-P |
|---|---|---|---|
| 114.09134 | L | | |
| 227.1754 | L | 1321.63 | 1223.6531 |
| 324.22817 | P | 1208.546 | 1110.5691 |
| 471.29658 | F | 1111.4932 | 1013.5163 |
| 542.3337 | A | 964.42479 | 866.44789 |
| 613.37081 | A | 893.38767 | 795.41078 |
| 684.40792 | A | 822.35056 | 724.37366 |

| | | | |
|---|---|---|---|
| 781.46069 | P | 751.31345 | 653.33655 |
| 878.51345 | P | 654.26068 | 556.28379 |
| 1045.5118 | S-Phospho | 557.20792 | 459.23102 |
| 1173.5704 | Q | 390.20956 | |
| 1260.6024 | S | 262.15098 | |
| | R | 175.11895 | |

Supplementary Table 4. Sense-specific qRT-PCR primers

| Primer name | Sequence (5’->3’) |
|---|---|
| RT_NP vRNA | AGCAAAAGCAGGGTAGATAATCAC |
| RT_NP mRNA | oligodT |
| RT_NP cRNA | AGTAGAAACAAGGGTATTTTTCTTTAATTGTCG |
| qPCR_NP vRNA Fwd | AGCAAAAGCAGGGTAGATAATCAC |
| qPCR_NP vRNA Rev | ATTTCAGTGGCATTCTGGCGTTC |
| qPCR_NP m/cRNA Fwd | GAGTCTTCGAGCTCTCGGACG |
| qPCR_NP m/cRNA Rev | GTCGTACTCCTCTGCATTGTCTCC |
| Human 18s-RT | TGGCAAATGCTTTCGCTCTGG |
| qPCR_18s Fwd | GCGCCGCTAGAGGTGAAATTC |
| qPCR_18s Rev | TGGCAAATGCTTTCGCTCTGG |

Supplementary Table 5. Primers to synthesize DNA-templates for v/c-NP77 RNA synthesis using in vitro transcription:

| Primer name | Sequence (5’->3’) |
|---|---|
| NP77 vRNA_IVT Rev | TAATACGACTCACTATAGAGTAGAAACAAGGGTATTTTTCTTTACGC |
| NP77 vRNA_IVT For | AGCAAAAGCAGGGTAGATAATCACTC |
| NP77 cRNA_IVT For | TAATACGACTCACTATAGAGCAAAAGCAGGGTAG |
| NP77 vRNA_IVT Rev | AGTAGAAACAAGGGTATTTTTC |
